# Stable chromosome configuration and loop-based polarization in animal symbionts

**DOI:** 10.1101/2023.12.21.572873

**Authors:** Tobias Viehboeck, Philipp M. Weber, Nicole Krause, Nelle Varoquaux, Frédéric Boccard, Ivan Junier, Silvia Bulgheresi, Virginia S. Lioy

**Affiliations:** Department of Functional and Evolutionary Ecology, Environmental Cell Biology Group, University of Vienna, Vienna, Djerassiplatz 1, A-1030 Vienna, Austria; Division of Microbial Ecology, Center for Microbiology and Environmental Systems Science, University of Vienna, Djerassiplatz 1, 1030 Vienna, Austria; University of Vienna, Vienna Doctoral School of Ecology and Evolution; Univ. Grenoble Alpes, CNRS, UMR 5525, VetAgro Sup, Grenoble INP, TIMC, 38000 Grenoble, France; Université Paris-Saclay, CEA, CNRS, Institute for Integrative Biology of the Cell (I2BC), 91198 Gif-sur-Yvette, France

## Abstract

Chromosome partitioning precedes the division of the cytoplasm, and its evolution is linked with the positioning of the division plane. So far, bacterial chromosome biology has heavily focused on transversally dividing, free-living ones. Here, we determined the chromosome organization of three longitudinally dividing *Neisseriaceae* exclusively inhabiting the oral cavity of mammals. We showed that in all three multicellular bacteria the origin of DNA replication is invariably located at the host-attached (proximal) pole. Next, 3C-seq revealed loop-based folding of the *ori* region in *Alysiella filiformis* and *Simonsiella muelleri*. Moreover, genes involved in cell motility, piliation and signal transduction mechanisms were specifically looped when transcriptionally and translationally active cells adhered to a substrate, but not when cultured in liquid. Overall, we propose that proximal positioning of the *ori* and loop-based folding of its surrounding DNA may mediate localized translation of proteins involved in host colonization.

## INTRODUCTION

In rod-shaped bacteria, chromosomes have been shown to adopt two main spatial configurations: left-*ori*-right (transverse) or *ori-ter* (longitudinal) ^1,2^. In the transverse configuration, the origin of DNA replication (*ori*) and the terminus of replication (*ter*) are at mid-cell, so that the left and right arms reside in opposite cell halves. In the longitudinal configuration, *ori* is located at one cell pole and *ter* at the opposite pole so that both left, and right chromosomal arms are parallel to the long axis of the cell. While model rods such as *Escherichia coli* and *Bacillus subtilis* can switch between transverse and longitudinal arrangements depending on the growth conditions or the cell cycle stage ^3–7^, the chromosome in *Caulobacter crescentus*, *Pseudomonas aeruginosa* and *Vibrio cholerae* maintains a longitudinal configuration throughout generations ^8–10^. In these three monoflagellated and vertically polarized bacteria, after the replication of the *ori*, the ParABS system actively segregates the duplicated *parS* regions towards the opposite pole ^11,12^. A variation on the theme of the longitudinal configuration is the so-called *ori-ter-ter-ori* configuration reported for the diploid *Actinomycetales Corynebacterium glutamicum* ^13^. In this configuration, the two *ori* are tethered to the poles in non-dividing cells, but both new-born *ori* are segregated to mid-cell, where they remain until fission is completed. Only then, they get tethered to the newly forming cell poles. While in all the aforementioned bacteria the intracellular localization of the *ori* can drastically vary during the cell cycle, in the longitudinally dividing *Candidatus* Thiosymbion oneisti the *ori* is maintained in the central third of the cell during its replication and lateral segregation, likely by the ParABS system ^14^. Taken together, irrespective of how the chromosome is configured, during replication and concomitant segregation, the subcellular localization of at least one of the sister *ori* along the long axis changes substantially in all bacteria studied so far ^15^, but the longitudinally dividing symbiont *Ca.* T. oneisti ^14^.

Besides displaying specific spatial dispositions and segregation modes, bacterial chromosomes may differ in their 3D internal organization, as revealed by chromosome conformation capture techniques coupled to deep sequencing (e.g., 3C-seq or Hi-C; ^7,16–21^. In all cases, Hi-C contact maps showed that chromosomes are segmented in sub-megabase chromosomal interaction domains (CIDs) whose boundaries often contain long and highly expressed gene clusters, indicating that intense transcription may limit the interactions between adjacent chromatin domains ^17,22^. Although CIDs are reminiscent of the eukaryotic topologically associated domains (TADs), they differ in their formation mechanisms. In mammalian cells, TADs are often delimited by chromosomal loops ^23^. These loops are observed at TADs boundaries as corner peaks and result from the structural maintenance of chromosomes (SMC) cohesin complexes and the CCCTC-binding factor (CTCF) binding the DNA at specific sites ^24–26^. In bacteria, chromosomal loops have been described in *B. subtilis.* Initially, some of these loops were proposed to be involved in the regulation of *ori* firing ^7,27^. More recently, loops were shown to be involved in some CID establishment during stationary phase via the action of the transcriptional factor Rok ^28^. Specifically, Rok was shown to form large nucleoprotein complexes that interact with each other over large distances, which eventually leads to anchored chromosomal loops ^28^. Finally, chromosomal loops have also been detected in euryarchaeota ^29^ and involved in the aggregation of high expressed insulated domains in the crenarchaeon *Aeropyrum pernix* ^30^. In the former case they are SMC-dependent but transcription independent, whereas in the latter, SMC-less organisms loops are transcription sensitive ^29,30^.

Despite the importance of chromosome orientation, segregation and conformation for cell functioning, only a limited number of bacteria have been studied so far ^12,19,31^. Here, we investigated the chromosomes of three multicellular *Neisseriaceae* commonly found attached to the oral mucosa of warm-blooded vertebrates ^32^: a canine strain of *Conchiformibius steedae*^33,34^, a caprine strain of *Alysiella filiformis* ^35,36^ and a human strain of *Simonsiella muelleri* ^36,37^ (Fig. 1a-c). By applying DNA fluorescence in situ hybridization on fixed cells (FISH), we showed that the chromosome configuration of these oral bacteria is stable, with the *ori* invariably localized at the host-attached (proximal) pole throughout their cell cycle, likely involving the centromeric protein ParB. Moreover, 3C-seq revealed the systematic presence of CIDs in exponential phase, most of them being delimited by long and highly expressed genes. Besides domains, we observed chromosomal loops in the host-proximal region of both *A. filiformis* and in *S. muelleri*, what we refer to as chromatin polarization hereafter. In *S. muelleri*, some loops delineated CIDs, similarly to mammal-like mechanisms associated with the formation of TADs. Strikingly, the abundance and the position of chromosomal loops were sensitive to the growing conditions (solid or liquid medium), as well as to transcription or translation inhibitors. Namely, growth in liquid medium led, in *A. filiformis,* to the enlargement of an *ori*-containing chromosomal loop and, in *S. muelleri,* to a ∼10-fold decrease in the number of chromosomal loops. Strikingly, in both oral bacteria, genes putatively involved in in motility and piliation were specifically looped when cells adhered to agar dishes, suggesting that loops might be associated with transertion phenomena involved in host colonization. Altogether, we propose that stable chromosome configuration and loop-based folding of chromosomes are cell biological mechanisms underlying host-bacterium interactions.

**Figure 1.**
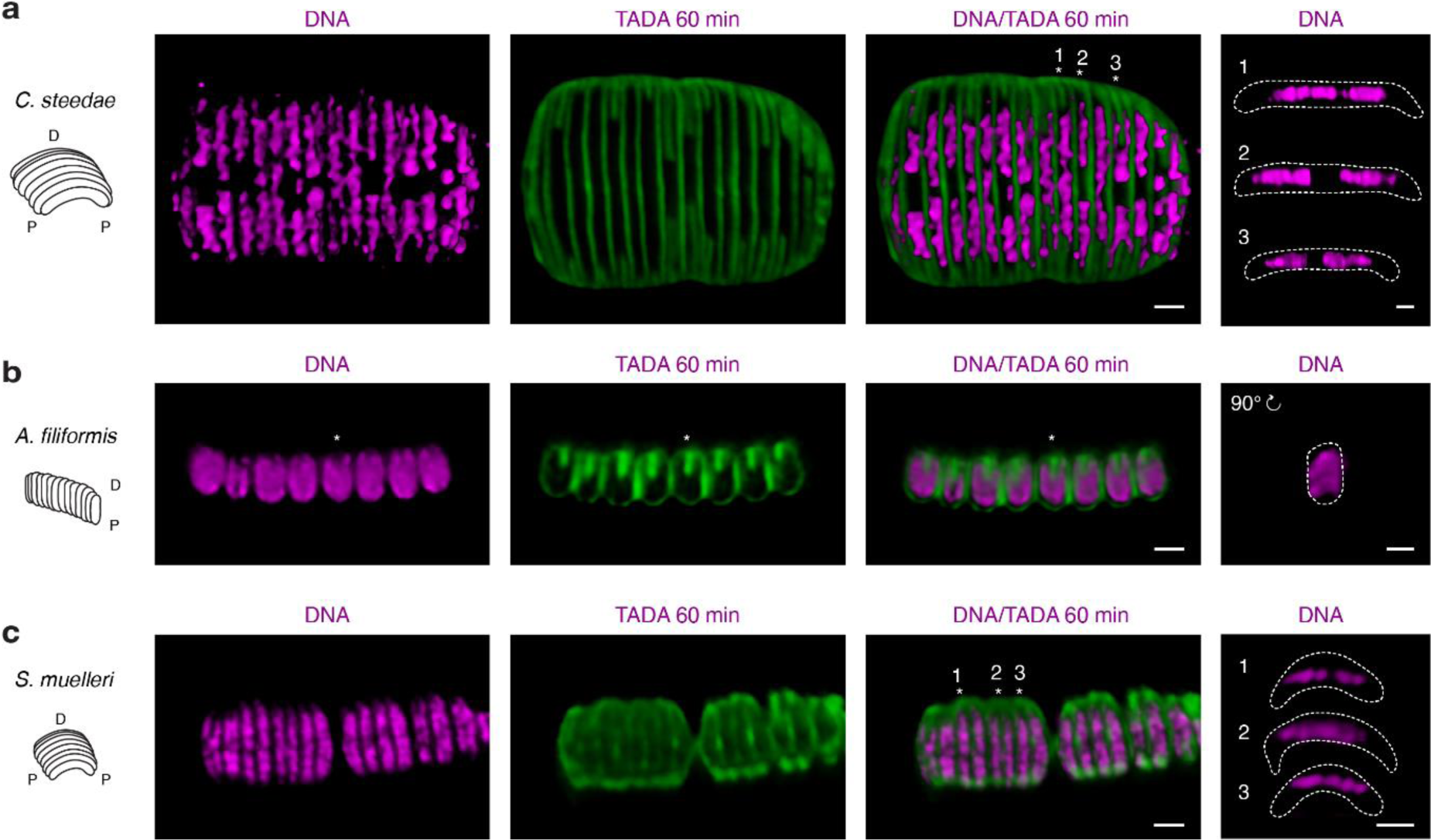
DNA localization in *C. steedae*, *A. filiformis* and *S. muelleri* cells. Schematic representations and confocal microscope images of *C. steedae* (a), *A. filiformis* (b) and *S. muelleri* labeled with the peptidoglycan precursor TADA for 60 min (green) and stained with Hoechst 33342 (magenta). (a) Viewed from the top, *C. steedae* DNA is either homogenously distributed (cell #1), excluded from midcell (cell #2) or X-shaped in cells undergoing late septation stages (cell #3). Rightmost panel displays a side view of cells 1-3. (b) Frontal view of an *A. filiformis* filament reveals a V-shaped DNA localization pattern. Rightmost panel displays a 90 degrees rotated view of the cells marked with an asterisk. (c) Viewed from the top*, S. muelleri* DNA is either excluded from midcell (cell #1), displaying a X-shaped pattern in cells undergoing late septation stages (cell #2) or homogenously distributed (cell #3). Rightmost panel displays a side view of cells 1-3. Scale bars are 1 µm.

## RESULTS

### Stable longitudinal chromosome configuration in three oral cavity symbionts

The three multicellular bacteria studied in this work are vertically polarized, as indicated by the host-proximal localization of their fimbriae ^32,37–39^. We will henceforth refer to the host-attached, fimbriae-rich pole of *A. filiformis* as the proximal pole and to the free pole as the distal pole. Regarding their division modes, *A. filiformis* divides in a distal-to-proximal fashion ^32^, whereas in the crescent-shaped *S. muelleri* and *C. steedae*, both poles are attached to the mouth epithelium which makes their mid-cell correspond to the region which is furthest away from the animal surface ^32,37^. Moreover, in *S. muelleri* and *C. steedae* the fimbriae localize to the side of the filament facing the oral mucosa, also referred to as the proximal (or ventral) side ^32,37^. Concerning their division mode, *S. muelleri* and *C. steedae* both septate from their host-attached poles toward mid-cell ^32^.

To determine their ploidy and chromosome configuration, we fixed exponentially growing *C. steedae, A. filiformis* and *S. muelleri* and subjected them to DNA fluorescence in situ hybridization (FISH) with specific probes specifically targeting their predicted *ori* and *ter* (Supplementary Fig. 1).

When *C. steedae* was subjected to DNA FISH, almost 50% of the cells (44%; n = 215) displayed two *ori* foci (Fig. 2a, Supplementary Table 1), and 76% (n = 236) displayed at least two *ter* foci (Fig. 2d, Supplementary Table 1), with *ori* invariably localized proximally (Fig. 2b-c), and the *ter* localized at mid-cell (Fig. 2e-f). Furthermore, lateral segregation of the *ori* was discernable (Supplementary Fig. 2a-c). As observed for *C. steedae*, most *S. muelleri* cells (47%; n = 188; Fig. 2g) had two *ori*, each at one pole (Fig. 2h-i). As for the *ter*, we detected one or two foci at mid-cell in most of the cells (61% and 39%, respectively; n = 293; Fig. 2j-l).

**Figure 2.**
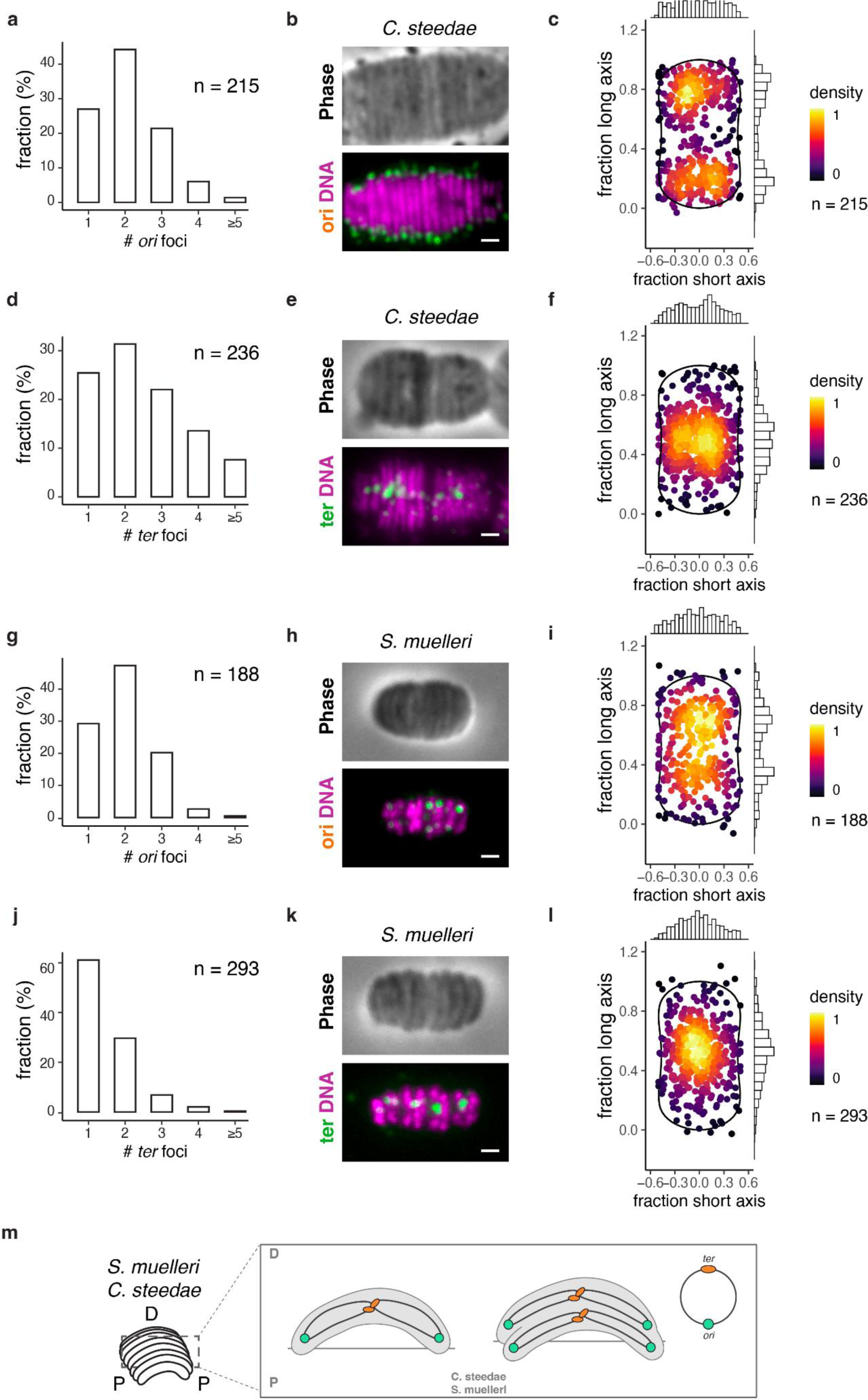
Stable *ori*-*ter*-*ter*-*ori* chromosome configuration in the diploid *C. steedae* and *S. muelleri*. (a, j) Histograms showing number of *ori* or (d,j) *ter* foci per cell. Confocal microscope images of representative filaments of *C. steedae* (b,e) and *S. muelleri* (h,k) subjected to either *ori* DNA FISH (b,h) or *ter* DNA FISH (e,k); phase contrast images are displayed in upper panels and corresponding FISH (green) and DNA (magenta) fluorescence images in bottom panels. (c,i) Subcellular localization of 215 *ori* foci in *C. steedae* and of 188 *ori* foci in *S. muelleri*. (f, j) Subcellular localization of 236 *ter* foci in *C. steedae* cells and of 293 *ter* foci in *S. muelleri*. (m) Model of *ori-ter-ter-ori* chromosome configuration and lateral chromosome segregation in the diploid crescent-shaped *S. muelleri* and *C. steedae*. Scale bars are 1 µm.

Multifork replication, as observed in fast-growing bacteria ^40^, could explain the relatively high *ori* count observed in *C. steedae* and *S. muelleri*. To exclude multifork replication in exponentially growing *C. steedae* and *S. muelleri*, we subjected them to marker frequency analysis (MFA), together with *A. filiformis* and *Escherichia coli.* We found that the *ori-ter* ratios of all three multicellular *Neisseriaceae* did not significantly change between exponentially growing and stationary phase cells, with values in the 1.3-1.9 range (Supplementary Fig. 3a-f). In *E. coli*, instead, the *ori-ter* ratio changed from 2.5 to 0.97 between exponentially growing and stationary phase cells (Supplementary Fig. 3g-h), as previously observed ^41^. This indicates that, contrary to *E. coli*, oral cavity symbionts do not undergo multifork replication and that *C. steedae* and *S. muelleri* are likely diploid.

Applied to *A. filiformis,* DNA FISH revealed that 98% of the cells (n = 379; Fig.3 a) have one *ori* and 96% (n=183; Fig. 3d) have one *ter*, indicating that this bacterium is likely monoploid. As for the configuration of the *A. filiformis* chromosome, we observed that, independently of the cell division stage*, ori* localized proximally (Fig. 3b-c and Supplementary Fig. 2d-e), whereas *ter* was distal (Fig. 3e-f and Supplementary Fig. 2f-g). Moreover, localization pattern analysis of *A. filiformis* containing either one or two *ori* foci (Supplementary Fig.2d-e) or either one or two *ter* foci (Supplementary Fig. 2f-g) indicated lateral chromosome segregation. Finally, after identification of the *A. filiformis* ParB protein using phylogenetic analysis (Supplementary Fig. 4a-b), we generated specific anti-ParB antibodies and used them for immunostaining (Supplementary Fig. 4c). The results were consistent with this centromeric protein mediating longitudinal chromosome configuration and segregation (Fig. 3 g-i).

**Figure 3.**
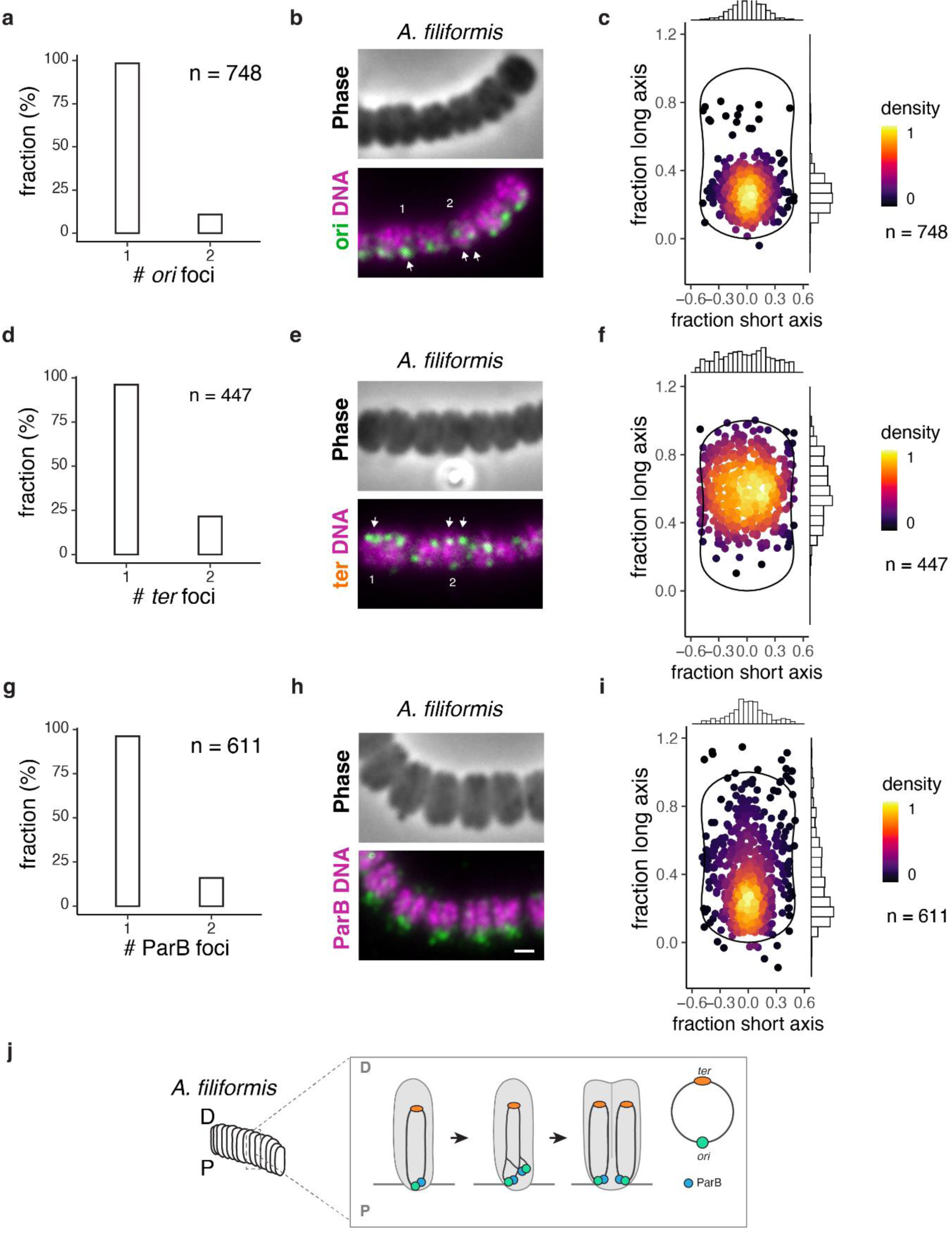
Stable *ori-ter* configuration in the monoploid *A. filiformis.* (a) Histograms showing number of *ori* foci (n = 748) and (d) *ter* foci (n = 447) per cell. (b,e) Confocal microscope images of representative filaments of *A. filiformis* subjected to *ori* DNA FISH (b) or *ter* DNA FISH (e); phase contrast images are displayed in upper panels and corresponding FISH (green) and Hoechst 33342 (magenta) fluorescence images in bottom panels. (c,f) Subcellular localization of 748 *ori* foci and 447 *ter* foci in *A. filiformis*. (g) Histogram showing number of ParB foci, (h) epifluorescence microscope image of an *A. filiformis* filament immunostained with anti-ParB antibody and (i) subcellular localization of 611 ParB foci. (j) Model of *ori-ter* chromosome configuration and lateral chromosome segregation in the monoploid *A. filiformis*. Scale bars are 1 µm.

Collectively, our results show that multicellular *Neisseriaceae*, irrespective of their ploidy and irrespective of their cell cycle stage, localize their *ori* proximally and their *ter* distally resulting in stable *ori-ter-ter-ori* (*C. steedae* and *S. muelleri,* Fig. 2m) or *ori*-*ter* (*A. filiformis,* Fig. 3j) configurations. Furthermore, based on its localization pattern, the centromeric protein ParB might mediate proximal positioning of the *ori* in *A. filiformis* (Fig.3 j). Finally, the finding of stable chromosome configurations in bacterial symbionts living attached to marine nematodes ^14^ (Supplementary Fig. 5) suggests that they might have convergently evolved to serve the symbiotic lifestyle.

### Starkly different chromosome architectures in three evolutionary related *Neisseriaceae*

Next, we used 3C-seq and RNA-seq to study the folding properties of the chromosomes of *C. steedae*, *A. filiformis*, and *S. muelleri* grown on solid medium and their putative relationships with gene expression. Despite the phylogenetic relatedness of these oral bacteria ^32^, genome-wide contact maps at a 5 kb-bin resolution revealed starkly different chromosome architectures (Fig. 4a-d). Firstly, only the maps of *C. steedae* and *S. muelleri* displayed a secondary diagonal, indicating the presence of low frequency, but significant, inter-arm DNA contacts (Supplementary Fig. 6a-b). This was consistent with the longitudinal organization of their chromosome (see previous section). By contrast, the contact map of *A. filiformis*, revealed rare inter-arm contacts despite the apparent longitudinally configuration of its chromosome and the presence of the condensin complex Smc-ScpAB and the ParABS system (Fig. 4b; Supplementary Fig. 6c).

**Figure 4.**
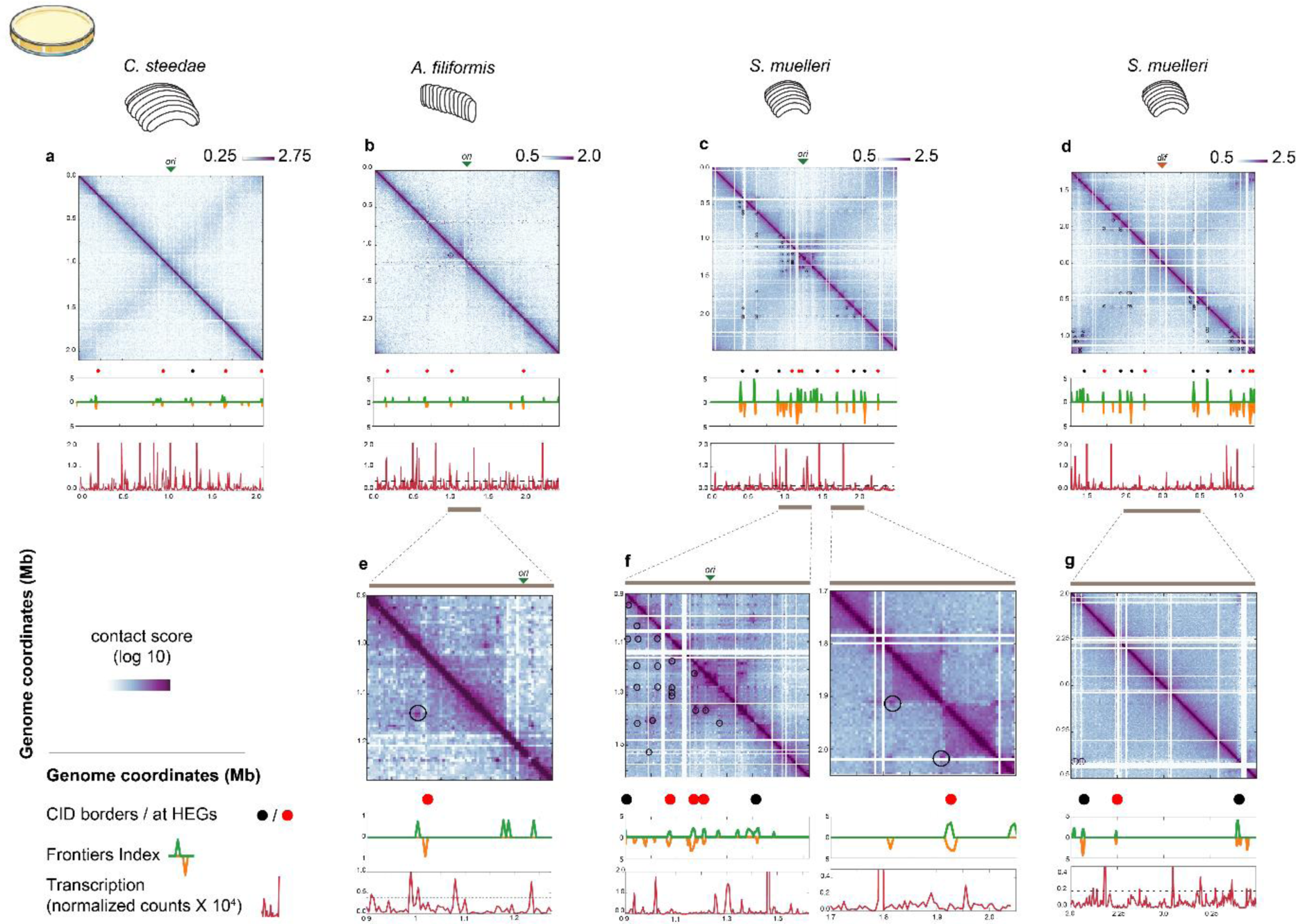
Normalized chromosome contact maps of *C. steedae*, *A. filiformis* and *S. muelleri* grown on solid medium. (a, b, c) *ori*-centered contact maps (green arrows denote *ori*), (d) *dif*-centered *S. muelleri* contact map (red arrows denote *dif*) of cells grown on solid medium, highlighting the loops observed in both replicates (black circles) in the lower half of each contact map. Below each map, the black or red circles show the position of highly expressed genes (HEGs) and rRNA operons if their presence correlates with a CID border. Below each track, the Frontier Index (FI) shows frontiers with upstream (green) or downstream (orange) bias of interaction and the normalized RNA-Seq counts in 5 kb bins show the transcriptomic data obtained in the same conditions that the contact map. The dashed line represents the threshold use to identify highly expressed regions. (e) Detail of (a) showing the *A. filiformis* loop right of *ori*. (f) Detail of (c) showing the *S. muelleri ori* region (left panel) and detail of (c) showing corner peaks of CIDs (right panel). (g) Detail of (d) showing the *S. muelleri* region around *dif*, with CID borders. The remaining replicates are shown in Supplementary Figs. 6-8.

Along the main diagonal, the 3C contact matrices revealed the presence of nested domains. To detect the chromosomal loci implicated in domains that span a wide range of sizes, we used the Frontier Index (FI), a tool that has been developed specifically to deal with the presence of multiple scales in the domain organization of 3C contact matrices ^42^. Specifically, the FI detects bins associated with either a significant upwards or a significant downwards interaction bias at any scale (up- or downstream peaks, respectively, Figure 4). In this context, we will henceforth refer to (1) loci corresponding to an up- or to a downstream peak as frontiers, (2) loci where an upstream peak is followed by a downstream peak (±3 bins) as domain boundaries and (3) loci between two domain boundaries as CIDs. In total, we identified 18, 16 and 37 frontiers in *C. steedae*, *A. filiformis*, and *S. muelleri*, respectively, with <50% of them corresponded to domain boundaries (Table 1, Fig. 4a-d; Supplementary Fig. 7a, Supplementary Fig. 8a-b and Supplementary Fig. 9a-b). In the case of substrate-attached *C. steedae* and *A. filiformis*, 80% (4 out of 5) and 100% (5 out of 5) of the domain boundaries matched long and/or highly expressed genes (HEGs, including *rrna* operons; Table 1; Supplementary Fig. 10a-b), consistent with previously observations in other bacteria ^7,13,16–18,22,43^. Notably, piliation genes (*pilMNOPQ*) located in the domain boundary situated at 1.7 Mb appeared strongly insulated from the rest of the chromosome (Fig. 4a). As for *S. muelleri*, the correlation between domain boundaries and HEGs was 45% in replicate 1 (only five out of the 11 boundaries matched HEGs), but was 69% in replicate 2 (Table 1, Supplementary Fig.10a-b).

**Table 1.** Number of frontiers, domain boundaries and co-occurrence with either highly expressed genes (HEGs) or chromosomal loops in different growing conditions.

| | Number of frontiers (up- or downstream peak) | Number of domain boundaries (overlapping up- and downstream peak ± 3 bins) | % of domain boundaries containing highly expressed genes (HEGs) | Number of detected chromosomal loops overlapping with frontiers | Number of detected chromosomal loops overlapping with domain boundaries |
| --- | --- | --- | --- | --- | --- |
| ***C. steedae*** |  |  |  |  |  |
| Solid medium | 18 | 5 | 80 | NA | NA |
| Liquid medium (exponential phase) | 20 | 12 | 92 | NA | NA |
| Liquid medium (stationary phase) | 13 | 4 | 100 | NA | NA |
| ***A. filiformis*** |  |  |  |  |  |
| Solid medium | 16 | 4 | 100 | 1 | 0 |
| Liquid medium (exponential phase) | 16*/25** | 7*/13** | 71*/62** | 1*/1** | 0 |
| Liquid medium (stationary phase) | 22 | 7 | 29 | 1 | 0 |
| Solid medium + Rifampicin | 20 | 5 | n.a. | 1 | 0 |
| Solid medium + Chloramphenicol | 15 | 5 | n.a. | 1 | 0 |
| ***S. muelleri*** |  |  |  |  |  |
| Solid medium | 32*/37** | 11*/16** | 45*/69** | 19*/17** | 12*/18** |
| Liquid medium (exponential phase) | 28 | 6 | 67 | 1 | 0 |
| Liquid medium (stationary phase) | 28 | 7 | 43 | 2 | 0 |
| **Solid medium**<br>**+ Rifampicin** | 18 | 2 | n.a. | 1 | 0 |
| **Solid medium**<br>**+**<br>**Chlorampheni**<br>**col** | 30 | 10 | n.a. | 1 | 4 |

\*Replicate 1; \*\*Replicate 2
HEGs: Highly expressed genes are genes that are found in regions in which the level of
transcription is classified as Cat\_4 (Supplementary Fig. 10 and see main text).

Collectively, we unveiled the large-scale organization of the chromosomes of three multicellular *Neisseriaceae*. Our results showed the absence of frequent inter-arm contacts in *A. filiformis* despite a longitudinal organization similar to *C. steedae and S. muelleri.* We also unveiled the presence of chromatin constrains that define multiple frontiers which are not linked to transcriptional activity in any of the studied symbionts. Finally, we have uncovered the existence of CIDs whose boundaries are not associated to HEGs, suggesting that intense transcription of long transcriptional units (or of a succession of them) is not the sole mechanism of chromatin organization in *S. muelleri*.

### Loop-based polarization of the chromosomes in substrate-attached *A. filiformis* and *S. muelleri*

Apart from domains, the contact maps of *A. filiformis* and *S. muelleri* revealed peaks of genetic interaction that we ascribed to chromosomal loops (indicated as black circles, Fig. 4c,e,f and Table 2). To identify their number unambiguously, we developed and applied a loop-detection algorithm to two contact maps established independently for each oral bacterium (replicate 1 and replicate 2). In the case of *A. filiformis*, we detected one 140 kb-long loop on the right side of the *ori* (Fig. 4b,e, Fig. 6a and Tables 1 and 2). Although this loop did not match any domain boundary (Tables 1 and 2), it resulted from the interaction of loci involved in host colonization (see last section of the Results).

**Figure 5.**
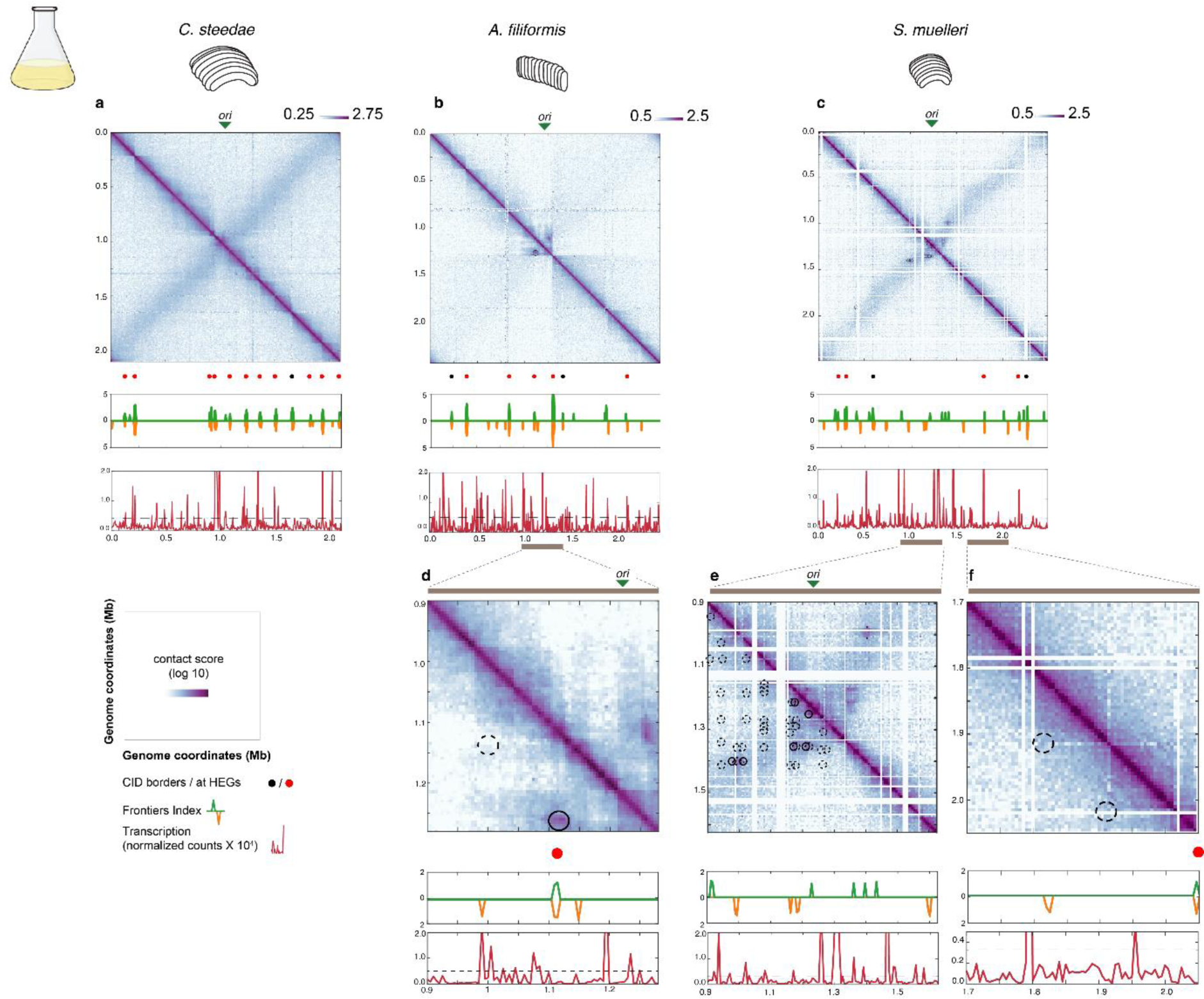
Normalized chromosome contact maps of *C. steedae*, *A. filiformis* and *S. muelleri* grown in liquid culture. (a, b, c) *ori*-centered contact maps (green arrows denote *ori*), highlighting the loops observed in both replicates (black circles) in the lower half of each contact map. Below each map, the black or red circles show the position of highly expressed genes (HEGs) and rRNA operons if their presence correlates with a CID border. Below each track, the Frontier Index (FI) shows frontiers with upstream (green) or downstream (orange) bias of interaction and the normalized RNA-Seq counts in 5 kb bins show the transcriptomic data obtained in the same conditions that the contact map. The dashed line represents the threshold use to identify highly expressed regions. (d) Detail of (b) showing the absence of the loop shown in Fig. 4e (dashed circle) and the emergence of a new loop containing the *ori* (closed circle). (e) Detail of (c) showing the absence of many loops (dashed circles) and the emergence of six loops (closed circles) in the vicinity of *S. muelleri* ori. (f) Detail of (c) showing the absence of the two corner peaks shown in Fig.4 f (right panel).

**Figure 6.**
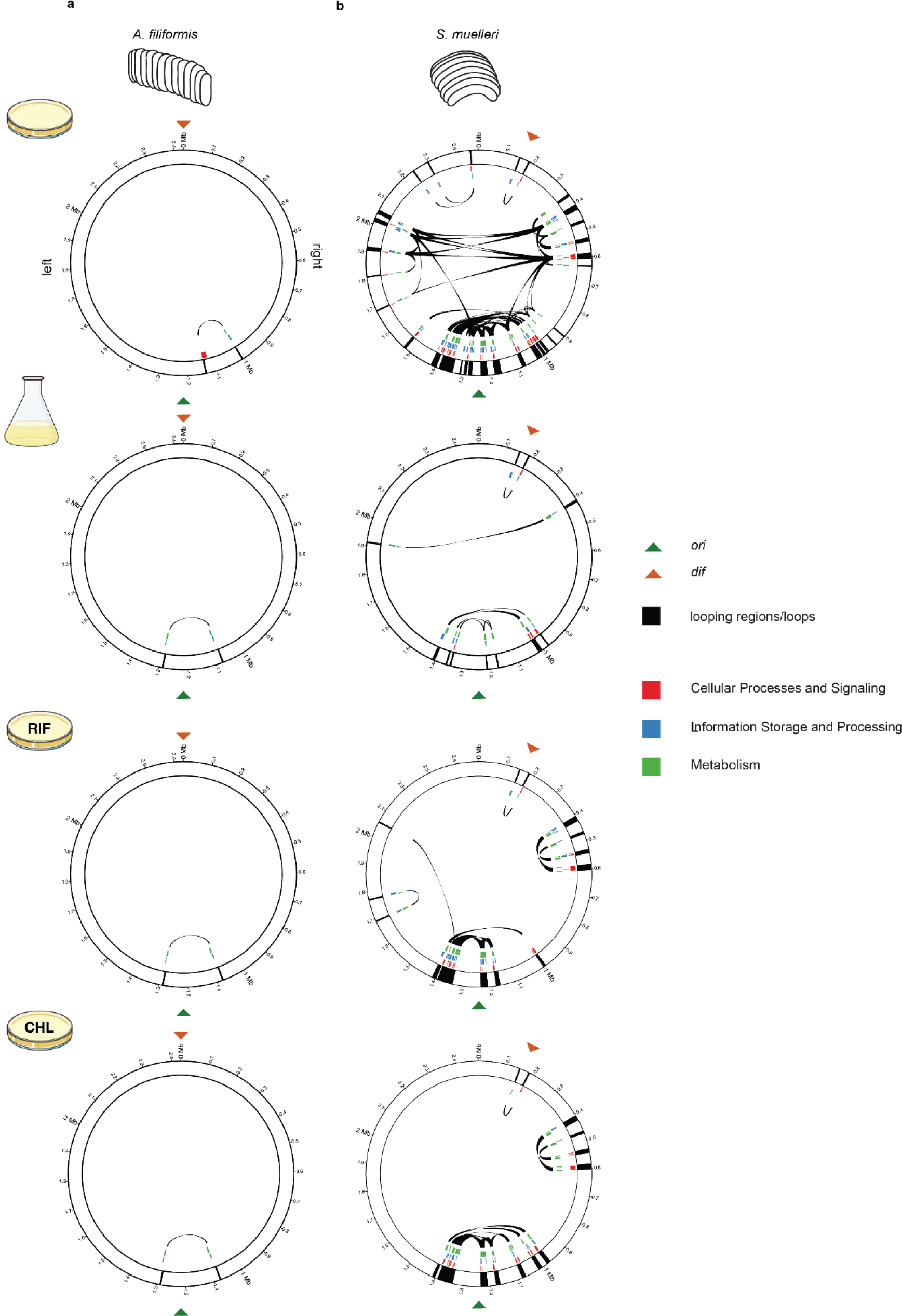
Circular representations of the *A. filiformis* and *S. muelleri* chromosomes and eggNOG-based annotation of genes contained in looping regions. (a, b) Circular representation of the *A. filiformis* and *S. muelleri* chromosomes when grown on solid medium (top panels) or in liquid medium (bottom panels), respectively. Looping regions are indicated as black lines on the white circle. The three inner colored circles represent the position of genes classified as one of the three COG categories ‘Cellular Processes and Signaling’ (red), ‘Information Storage and Processing’ (blue), and ‘Metabolism’ (green). Connections between looping regions (loops) are shown in black. Green arrows indicate the position of *ori*, and the red arrows indicate the position of *dif*.

**Table 2.**
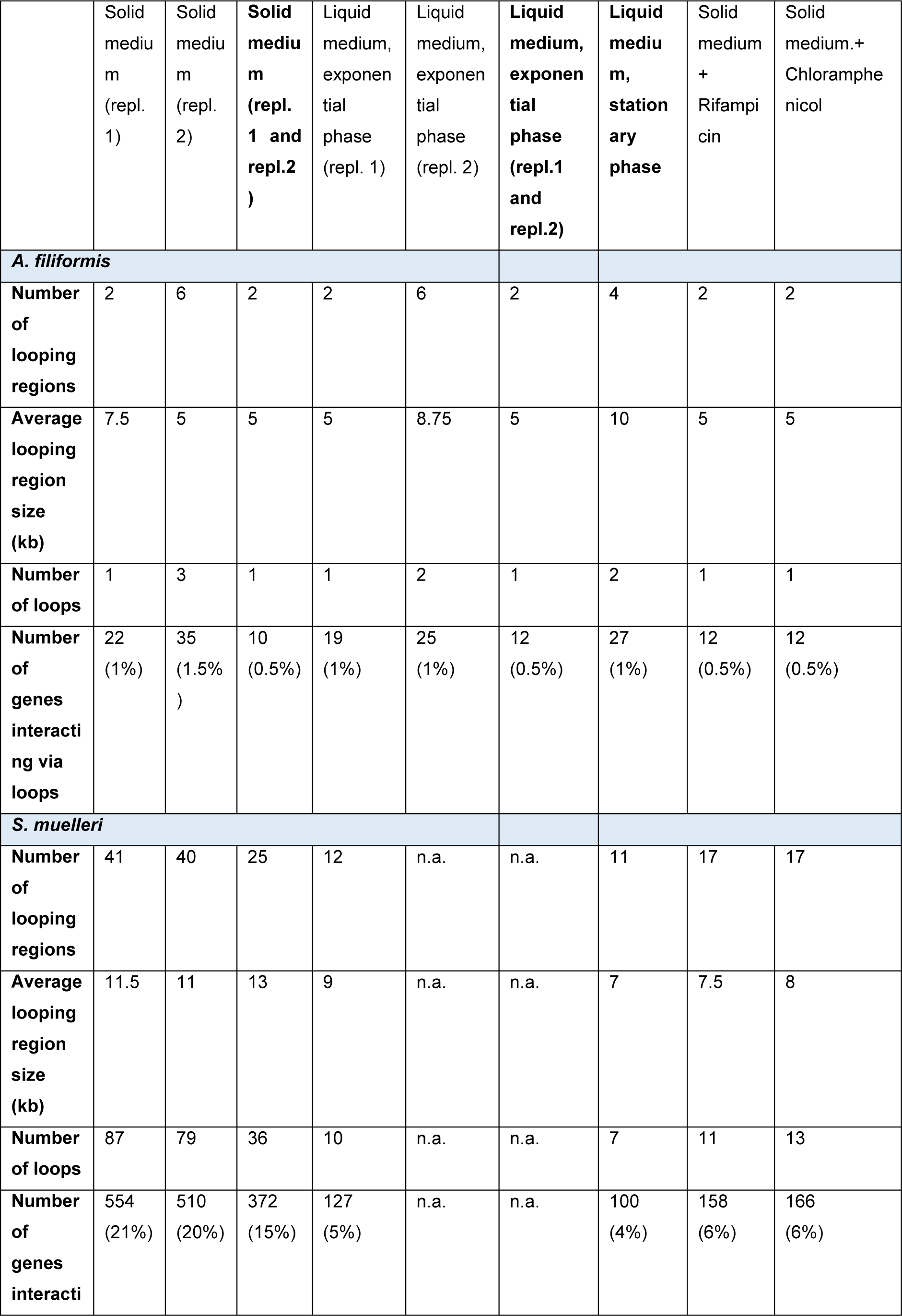
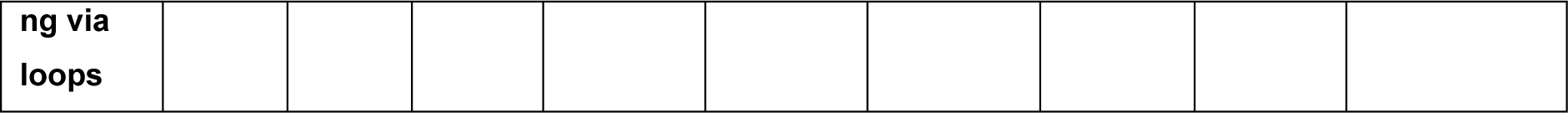
Number of looping regions, of loops and of genes interacting via loops in *A. filiformis and S. muelleri*.

In the case of *S. muelleri*, our loop-detection algorithm revealed 36 distinct loops (Fig. 4c-d, black circles) engaging 25 different chromosomal regions (referred to as looping regions hereafter), with each looping region being connected to 2.4 other looping regions on average. These loops were not distributed homogenously throughout the chromosome. Instead, they generated a highly condensed region around *ori* (between positions 0.8 and 1.4 Mb), a second one on its right side (between positions 0.3 and 0.7 Mb) and a third one on its left side (between 1.7 and 2.1 Mb) (Fig. 4c-d,f, Fig. 6b), leaving a ≍1.4 Mb loopless region around *dif*. Of note, 50% of the loops matched domain boundaries (Fig. 4c-d, f and Tables 1 and 2).

Altogether, contact maps of the human symbiont *S. muelleri* unveiled a loop-based organization of its chromatin resembling that seen in mammalian cells. Furthermore, the identified loop in *A. filiformis* close to *ori* together with the strong bias of *S. muelleri* loops for the proximal (*ori*-containing) region imply that the chromosomes of these bacteria are not homogeneously folded along the cell long axis. We will hereafter refer to this phenomenon as loop-based polarization of the *A. filiformis* and *S. muelleri* chromosomes.

### Loss of substrate attachment affects loop-based chromosome folding

To assess whether losing the attachment to a substrate would result in a depolarization of chromosome folding (i.e., in a loss of chromosomal loops), we performed 3C-seq in *A. filiformis* and *S. muelleri* grown in liquid culture and fixed either in exponential or stationary phase (Fig. 5b-f, Supplementary Fig. 8c-e, Supplementary Fig. 9c-e). Furthermore, we also performed 3C-seq on liquid-grown *C. steedae* (Fig. 5a) to know whether changes in growing conditions or growth phase would affect the number of frontiers.

Overall, the global 3D organization of the chromosomes of liquid grown *C. steedae*, *A. filiformis* and *S. muelleri* was conserved, e.g., we observed a secondary diagonal in *C. steedae* and *S. muelleri*, and rare inter-arm contacts in *A. filiformis.* In addition, FI detected a similar number of frontiers in the three symbionts (Fig. 5a-c, Supplementary Fig. 7b, Supplementary Fig. 8c-d and Supplementary Fig. 9c-d), albeit exponential phase *C. steedae* and liquid-grown *A. filiformis* (irrespective of the growth phase), displayed more domain boundaries compared to bacteria grown on agar dishes (Table 1). Also of note, more domain boundaries corresponded to HEGs in liquid-grown *C. steedae*.

Most strikingly, in liquid growing *S. muelleri* the number of chromosomal loops decreased from 36 to seven irrespective of the growth phase (Fig. 5c, e-f, Supplementary Fig.9c-e and Table 2). Moreover, only one of the seven loops detected in liquid-grown *S. muelleri* matched a frontier (Table 1). Concerning *A. filiformis*, the 140 kb-long *ori*-containing loop detected in solid-grown cells (Fig. 4b,e and Fig. 6a top panel) became 215 kb-long in cells grown in liquid culture, irrespective of the growth phase (Fig. 5b,d, Fig. 6a bottom panel and Supplementary Fig. 8c-e).

All in all, loss of substrate attachment mainly affected the loop-based organization of the chromatin in *A. filiformis* and *S. muelleri*. Namely, loss of substrate attachment led to the formation of a longer, *ori-*containing chromosomal loop in *A. filiformis* and to an 80% loss of chromosomal loops in *S. muelleri*, with only seven left in the immediate vicinity of its *ori*.

### Loop-based polarization of the chromosomes in substrate-attached symbionts may facilitate localized translation of proteins involved in host colonization

To assess the function of chromosomal loops, we first tested the transcriptional levels of the genes they involved in substrate-attached *A. filiformis* and *S. muelleri* (Supplementary Fig.10c; Supplementary Data Set 1). In *A. filiformis*, we did not find any significant difference in the median level of transcription of looped genes versus the median level of transcription of all the genes. In substrate-attached *S. muelleri*, however, transcripts of looped genes were scarce (p<0.01), except for those encoding for translation, ribosomal structure and ribosomal biogenesis proteins (p<0.05) (Supplementary Data Set 1). Of note, growth of *S. muelleri* in liquid medium did not increase the abundance of transcripts corresponding to looped genes suggesting that looping was not responsible of the low abundance of their transcripts in substrate-attached cells.

In our quest to understand the biological function of chromosomal loops, we then resorted to automatic annotation of the looping regions (based on the eggNOG database). In substrate-attached *A. filiformis*, the two looping regions contained genes putatively needed for colonization of the host, including a signal recognition particle (SRP)-containing,15 kb-long adhesin and a transcription repressor involved in efflux mediated antimicrobial resistance (Fig. 6a top, Supplementary Tables 5 and 6 and Supplementary Data Set 1); ^44^. However, when *A. filiformis* was grown in liquid medium, none of the looped genes belonged to the COG category *Host adhesion and/or immune evasion*. Instead, looped genes belonged to either *Information storage and processing* or to *Metabolism* (Fig. 6a bottom, Supplementary Fig. 11a and Supplementary Data Set 1).

We then annotated the genes situated in the looping regions in *S. muelleri* growing on solid medium using the eggNOG database (Fig. 6b, Supplementary Fig. 11a and Supplementary Data Set 1). Firstly, we found the following solid medium-specific COG categories: *Cell motility* (including *pilA* and a gene encoding for a prepilin-related protein), *Defense mechanisms* (including the antibiotic resistance gene *ampD*) and *Signal transduction mechanisms*. Of note, 40% of cell motility genes were looped when *S. mulleri* was grown on agar plates. Concerning the “second most-looped” COG category, *Intracellular trafficking/Secretion*, although this category is not solid substrate-specific, the Type 4 Secretion System genes virB3-6 and the gene encoding for the SRP receptor *ftsY* were only looped when cells were substrate-attached. Also noteworthy were the COG categories *Cell wall/membrane/envelope biogenesis* and *Coenzyme transport and metabolism*. Although these two categories were not solid medium-specific, *wecA* and *neuABC* genes (encoding for capsule or lipopolysaccharide sialyating enzymes), two spermidine synthases and an adhesin were only looped in substrate-attached *S. muelleri* (Fig. 6b, Supplementary Fig. 11a and Supplementary Data Set 1). Finally, among the genes which could not be classified into any COG category, but that were only looped in substrate-attached *S. muelleri* and that may be involved in host adhesion and/or immune evasion, we found three genes encoding for immunoglobulin-binding proteins (*tspB*) and three copies of a gene encoding for zonula occludens toxins (*zot*) (Fig. 6b, Supplementary Fig. 11a and Supplementary Data Set 1).

In summary, functional annotation of highly frequently interacting (looped) genes suggests that, at least some of the loops may result from transertion, i.e., from DNA-membrane interaction due to the coupled transcription and translation of membrane or secreted proteins associated with host colonization. Consistent with this hypothesis, blocking transcription or translation in substrate-attached *A. filiformis* had the same effect as growing them in liquid culture (Table 1, Fig. 6a, Supplementary Fig.12). Along the same line, blocking translation in substrate-attached *S. muelleri* led to the loss of loops engaging genes encoding for cell motility proteins, pili, the SRP receptor and hemolysin (Table 1, Fig. 6b, Supplementary Fig.11 and Supplementary Fig.12).

## DISCUSSION

During the past decade, studies on symbiont cell biology revealed that rod-shaped bacteria may specifically localize their fimbriae to their host-attached poles ^39^. Moreover, bacterial symbionts orient their cytokinetic machineries orthogonal to the animal surface and septate longitudinally, so that both daughter cells stay attached to the host by one pole ^39,45,46^. Recently, we found that, thanks to a bidimensional segregation mode, a longitudinally dividing symbiont could keep its chromosome orientation toward its host throughout generations ^14^. Here, we expanded on the capacity of symbiotic cells to polarize their components by focusing on the spatial disposition and architecture of their genetic material.

Firstly, we found that all the symbionts studied here configure their chromosomes longitudinally throughout the cell cycle, i.e., the *ori* stays proximal irrespective of the ploidy and of chromosome segregation stage. This is achieved by lateral chromosome segregation, likely active and ParABS-mediated. Indeed, in *A. filiformis*, we identified seven *ori*-adjoining *parS* sites, and ParB signal was proximal throughout the cell cycle. Similarly, in the rod-shaped and nematode symbiont *Candidatus* Thiosymbion hypermnestrae, ParB foci are proximal at all cell cycle stages, but the very last one (Supplementary Fig.5). As for the nematode symbiont *parS* sites, we identified two of them in the vicinity of the *ori*. Therefore, in *A. filiformis* and *Ca.* T. hypermnestrae, ParA might drive lateral segregation of *ori*-ParB. As for *C. steedae* and *S. muelleri*, the former has two *parS* sites in the vicinity of the *ori* and four in the vicinity of the *ter*, whereas the latter has six *ori*- and two *ter*-adjoining *parS* sites. Further, in *C. steedae* and *S. muelleri*, 3C contact maps revealed the presence of inter-arm contacts (Supplementary Fig.6), suggesting that ParB mediates the loading of Smc-ScpAB at the *parS* sites situated in the vicinity of the *ori*. Interestingly, *B. subtilis* harbors eight *parS* sites in the *ori* region and each of them presents a specific ParB and Smc-ScpAB recruitment efficiency ^47^. Characterization of the affinity of ParB for the different *parS* sites is necessary to determine whether ParB has a higher affinity for *parS* sites situated in the vicinity of the *ori* than for the distal ones, and whether this may play a role in differential recruitment of the Smc-ScpAB complex in *S. muelleri* and *C. steedae*.

Secondly, DNA FISH-based count of *ori* and *ter* foci and MFA indicate that *S. muelleri* and *C. steedae* are likely diploid, i.e., newborn cells have two *ori* and two *ter* and therefore two copies of the same chromosome. Although about one third of *S. muelleri* or *C. steedae* displayed only one *ori* focus (instead of two), we ascribe this to an insufficient detection efficiency of the DNA FISH technique. Likely for the same reason, most *S. muelleri* cells displayed only one *ter* focus instead of two foci. Diploidy has also been reported in pathogenic *Neisseriaceae* ^48^, in the actinomycetes *C. glutamicum* ^13^ and in the Euryarchaeon *Methanothermobacter thermautotrophicus* ^49^. Because bearing two (or more) chromosome copies increases the chance of correctly repairing the DNA by homologous recombination, we hypothesize that *C. steedae* and *S. muelleri* evolved diploidy to withstand stress ^50^ including that exerted by bacteriophages or transposable elements ^51^. In this regard, the identification of prophage gene clusters and three transposase-encoding genes is noteworthy (Supplementary Fig. 9b). Alternatively, or additionally, diploidy may have evolved to maintain the intracellular position of genetic loci in longitudinally dividing curved bacteria, which adhere to the animal surface with both poles.

A third point is the spatial disposition of the genetic material inside the cell. Up to this work, all studied bacterial chromosomes displayed highly variable chromosome configurations: either *ori*, *ter* or both drastically changed their intracellular localization pattern during DNA replication and segregation (or when experiencing different growth phases) ^1,31^, except for the longitudinally dividing *Ca.* T. oneisti, where the *ori* is kept at mid-cell throughout generations and irrespective of the cell cycle stage ^14^. We currently do not know what pushed the oral symbionts to evolve formidably stable chromosome configurations. One evolutionary driver could have been the necessity to keep the genes where their products are needed, especially if these products are essential for the symbiont survival. Apart from possessing a genetic repertoire typical of aerobic chemoorganoheterotrophs, our annotations indicate that the oral cavity symbionts cannot synthesize vitamin B_12_ and, intriguingly, two genes encoding for transporters mediating its uptake (the TonB-dependent-vitamin B_12_ transporter BtuB and a vitamin B_1_ transporter) are situated close to the *ori* in *S. muelleri* and are therefore, thanks to the stable chromosome configuration, invariably proximal (Supplementary Data Set 1). The hypothesis that stable chromosome configurations evolved to facilitate the intracellular localization of membrane or secreted proteins where they are most needed relies on the assumption that mRNAs may be kept where they are transcribed, as shown for *C. crescentus* ^52^. An additional piece of data linking the intracellular localization of bacterial genes to the place they have to function is that translation of membrane proteins may trigger chromosomal locus repositioning ^53,54^. Intriguingly, incubation in 25 µg/ml chloramphenicol led to nucleoid compaction in both *A. filiformis* and *S. muelleri* (data not shown). However, localization pattern analysis of selected *A. filiformis* and *S. muelleri* mRNAs is needed to find out whether its chromosome serves as a template for the proximal localization of mRNAs involved, for example, in the uptake of essential nutrients from the host (e.g., the mRNA encoding for the vitamin transporter Btub).

Finally, in the case of *C. steedae* and *S. muelleri* (adhering to a solid substrate or planktonic) a secondary diagonal was visible in the contact maps consistent with the stable longitudinal organization revealed by DNA FISH. However, a secondary diagonal does not seem to be a prerequisite of chromosome longitudinal organization, as we detected rare inter-arm contacts in *A. filiformis* which also has an *ori-ter* configured chromosome. This suggests that in this monoploid symbiont the two chromosome arms are laying longitudinally, but further away from each other than in the thinner, longer and apparently diploid *C. steedae* and *S. muelleri*. Alternatively, the lack of inter-arm contacts in *A. filiformis* might be due to the absence of an SMC-based, *ori-ter* oriented loop extrusion mechanism ^55^.

Prior to this study, intensively transcribed loci were found in at least 60% of the CID boundaries ^7,13,17–19,22,43^. The correlation between domain boundaries and HEGs is consistent with the previous findings except for stationary phase *A. filiformis* and *S. muelleri* (Table 1), possibly because the number of frontiers was flawed by the intrinsic noise associated to 3C matrices (Supplementary Figs. 7-9). Furthermore, in these symbionts, we observed biased interactions in a single direction around certain loci, meaning that these loci do not separate two consecutive CIDs. Interestingly, these frontiers are not associated with HEGs, but with chromatin constrains that remain to be identified.

Concerning chromosomal loops, in substrate-attached *S. muelleri* that were actively transcribing and translating, they were frequently found at domain boundaries (Tables 1 and 2), resembling the corner peaks demarcating mammalian TADs. Their association with TADs can be explained by the following model: the cohesin complex binds the chromatin at a random place, extrudes DNA bi-directionally through a loop and is eventually blocked by the CCCTC-binding factor CTCF bound at the borders of the TAD ^56^. The blocking indeed promotes the stabilization of the loop, which thus manifests as a corner peak in the contact map ^26,57^. In *B. subtilis*, the transcriptional factor Rok, likely assisted by the SMC complex, is associated to chromosomal loop formation delineating CIDs in stationary phase ^28^. Whether a similar system is responsible for chromosomal looping and domain formation in substrate-attached *S. muelleri* remains to be demonstrate.

Functional annotation suggested that, at least some *S. muelleri* chromosomal loops, may result from localized transcription and translation of proteins involved in cellular motility, piliation and immune evasion. Localized transcription and translation of host colonization factors, facilitated by stable chromosome configurations, would allow, when cells are attached to the oral mucosa, to efficiently translate and assemble (or transert) membrane-bound molecular complexes required for host colonization. Consistent with this hypothesis, in *A. filiformis*, an adhesin gene and a gene involved in efflux-mediated antibiotic resistance were looped only when actively transcribing and translating cells adhered to a substrate, suggesting that, like in *S. muelleri*, chromosomal loops may be involved in host adhesion and/or immune evasion. (Fig. 6b, Supplementary Tables 5 and 6 and Supplementary Data Set 1). Along this line, transertion of the type 3 secretion system was recently shown in *Vibrio parahaemolyticus* incubated with bile acids ^54^.

Given that 3C-seq did not reveal any chromosomal loop in *C. steedae* (Fig. 4a, Fig. 5a and Supplementary Fig. 7), irrespective of the culture conditions, chromosomal loops do not seem a *sine qua non* feature of bacteria thriving on animal surfaces. Instead, loop-based chromosome polarization might be an exclusive feature of multicellular *Neisseriaceae* belonging to the M1 lineage (*Alysiella* and *Simonsiella* spp.) ^32^. Although chromosome conformation capture did not reveal chromosomal loops in the unicellular *Neisseriaceae Neisseria elongata* (data not shown), we must study more *Neisseriaceae* species and isolates to test this hypothesis.

In conclusion, it is possible that stable chromosome configurations, together with the presence of chromosomal loops in *A. filiformis* and *S. muelleri*, mediate localized (proximal) translation of proteins involved in host colonization (e.g., pili, enzymes for extracellular polysialic acid synthesis, adhesins, zonula occludens toxins, immunoglobulin-binding proteins). More generally, we propose that specific chromosome positioning and folding may represent cellular adaptations to an obligate symbiotic lifestyle.

## AUTHOR CONTRIBUTIONS

T.V. did 3C experiments, marker frequency analysis, RNA sequencing, genome sequencing and assembly, bioinformatic and formal analyses, visualization, wrote and revised the manuscript. P.M.W. did DNA staining, DNA FISH, formal analyses and visualization, wrote and revised the manuscript; N.K. did DNA staining, DNA FISH and formal analyses. I.J. performed computational analyses of 3C-seq data, developed the loop detection algorithm and revised the manuscript. N.V. performed preliminary FI analysis. B.D. did DNA FISH. F.B. acquired funding. V.S.L. performed preliminary 3C analysis, conceptualized and supervised the work, acquired funding, provided resources, wrote, and revised the manuscript. S.B. conceptualized and supervised the work, acquired funding, provided resources, wrote, and revised the manuscript.

## ACKNOWLEDGEMENTS

This work was supported by the Austrian Science Fund (FWF) project P28593 (T.V. P.M.W, N.K.), FWF project P28743 (T.V., P.M.W), FWF doc.funds MAINTAIN (T.V.), a DOC-fellowship from the Austrian Academy of Science (P.M.W.), and a PhD completion grant from the University of Vienna (P.M.W.), by the French National Research Agency: grant number ANR-20-CE35-005 (V.S.L), by the 80 Prime CNRS project MIMIC (F.B. and I.J.) and SIRIG (N.V and V.S.L.) through the MITI interdisciplinary exploratory research porgramms. We thank Bocar Diallo for performing some DNA FISH experiments, to Frédéric Veyrier (INRS, Quebec, CA) for introducing *Conchiformibius steedae* to us and for sharing the RNAseq protocol, the members of the F.B. and S.B. laboratories for fruitful discussions and to S. Bury-Moné for careful reading and feedback on the article. We acknowledge the sequencing and bioinformatics expertise of the I2BC High-throughput sequencing facility, supported by France Génomique (ANR-10-INBS-09). We are also indebted to the staff of the VBCF NGS Unit (Laura-Maria Bayer) for assistance with Oxford Nanopore MinION sequencing. The computational results of this work have been achieved using the Life Science Compute Cluster (LiSC) of the University of Vienna.

## LEAD CONTACT AND MATERIALS AVAILABILITY

Further information and requests for resources and reagents should be directed to and will be fulfilled by the lead contact, Silvia Bulgheresi. This study did not generate new strains or unique reagents.

## DATA AND CODE AVAILABILITY

Data for genome sequencing has been deposited in Bioproject PRJNA687213 (*A. filiformis*), PRJNA687218 (*S. muelleri*), and PRJNA779076 (*C. steedae*), and genomes have been deposited in GenBank as GCF_020162295.1 (*A. filiformis*), GCF_020162275.1 (*S. muelleri*), and GCF_023547005.1 (*C. steedae*). Data for marker frequency analysis has been deposited in Bioproject PRJNA795449 (*A. filiformis*), PRJNA795451 (*S. muelleri*), and PRJNA795450 (*C. steedae*), and corresponding non-normalized, 1kb-binned read count files in Gene Expression Omnibus (GEO) GSE194131 (*A. filiformis*), GSE194134 (*S. muelleri*), and GSE205139 (*C. steedae*). Data for RNA-Seq has been deposited in Bioproject PRJNA795348 (*A. filiformis*) and PRJNA795350 (*S. muelleri*), and corresponding raw read count files in GEO GSE194132 (*A. filiformis*), and GSE194135 (*S. muelleri*). Data for 3C-seq has been deposited in Bioproject PRJNA837061 (*A. filiformis*), PRJNA837062 (*S. muelleri*), PRJNA1052629 (*C. steedae),* and corresponding matrices in GEO GSE250400 (*A. filiformis*), GSE250402 (*S. muelleri*), and GSE250401 (*C. steedae*). SRA accession numbers for sequencing reads and GEO accession numbers for data files are listed in Supplementary Table 4. The documentation for the ImageJ plugin Fil-Tracer can be accessed here: https://sils.fnwi.uva.nl/bcb/objectj/examples/Fil-Tracer/MD/Fil-Tracer.html.

## EXPERIMENTAL MODEL AND SUBJECT DETAILS

The bacterial strains *Alysiella filiformis* (DSM 16848), *Simonsiella muelleri* (DSM 2579) and *Conchiformibius steedae* (DSM 2580) have been received from the German Collection of Microorganisms and Cell Cultures GmbH (DSMZ). All *Neisseriaceae* strains were prepared for experimentation as described below unless otherwise noted. *A. filiformis* cultures were pre-cultured by inoculating liquid peptone-yeast (PY) medium with glycerol stocks stored at −70°C and grown overnight at 37°C at 120 rpm, transferred to fresh PY medium the next day and grown until the specified OD_600_ (see each section). *S. muelleri* and *C. steedae* cultures were streaked out from glycerol stocks onto BSTSY agar plates, grown overnight at 37°C, and colonies transferred into liquid BSTSY medium, and grown at 37°C at 120 rpm until the specified OD_600_ (see each section).

For sampling symbiotic nematodes, sediment samples were collected on multiple field trips (2015-2019) in 1 m depth from a sand bar off Carrie Bow Cay, Belize (16°48’11.01’’N, 88°4’54.42’’W). Specimens of *Robbea hypermnestrae* were extracted from the sediment by stirring the sand in seawater and pouring the supernatant through a 63 µm mesh sieve. The retained material was transferred into a Petri dish and single nematodes were handpicked using pipettes under a dissecting microscope. For DNA FISH, whole symbiotic nematodes were fixed in 3 or 4% PFA for 12-14 h at 4°C, washed with 70% ethanol and stored in 70% ethanol at −20°C. Fixed symbiotic nematodes were transported from Carrie Bow Cay to the University of Vienna deep-frozen.

## METHOD DETAILS

### Generation of DNA fluorescence in situ hybridization (FISH) probes

We used the genome draft of *Ca*. T. hypermnestrae (Genbank accession number GCA_020443945.1) to design specific primers against the *Ca*. T. hypermnestrae *ori* and *ter* regions, as well as against DNA stretches containing the *parS2* (on a different contig than *ori* and *parS1)* ^58^ or the *ftsQAZ* operon (Supplementary Table 1; *Ca*. T. hypermnestrae *dif* site sequence: ACTTTACATAATATACATTATGCGTAA) ^59^. *Robbea hypermnestrae* nematodes were rehydrated in phosphate-buffered saline (PBS) and bacteria were detached from the worms by sonication. Subsequently, 1 µl of bacterial suspension was used as template in each 25 µl PCR reaction (primer sequences and PCR conditions are listed in Supplementary Tables 2 and 3). A 2,858 nt-long fragment containing the *dnaA* and *dnaN* genes (targeting the *ori*), a 3,690 nt-long fragment containing the *ftsQAZ* operon, a 3,524 nt-long fragment containing the *parS*2 site (targeting an *ori*-proximal site) and a 3,239 nt-fragment containing the predicted *dif* site (targeting a *ter*-proximal site) were amplified. Each purified fragment was then used as template to PCR-amplify dsDNA polynucleotide probes (referred to as *ori*, *ftsQAZ*, *parS2* and *ter*, respectively) (see Supplementary Tables 2 and 3 for primers and PCR conditions).

For the oral cavity symbionts *Alysiella filiformis* and *Simonsiella muelleri*, we used Gene-PROBER (Moraru, 2021) to design specific primers hybridizing with the origin or terminus of DNA replication or in their immediate vicinity (Supplementary Fig.2). For gDNA extraction, cells were lysed using lysozyme and Proteinase K, DNA extracted using Phenol-Chloroform and precipitated using ammonium acetate (2.5 M final concentration), two volumes of 100% ice-cold Ethanol, and 5 µg/ml glycogen. RNA was degraded by incubation with 10 mg/ml RNase A for 30 min at 37°C, and DNA cleaned up using DNA Clean & Concentrator-5 (Zymo). Extracted gDNA was used as a template in each 25 µl PCR reaction (primers sequences and PCR conditions are listed in Supplementary Tables 2 and 3), amplifying a 9,232 nt-long *ori* fragment and a 6,778 nt-long *ter* fragment from *A. filiformis*, an 8,158 nt-long *ori* and an 8,187 nt-long *ter* fragment from *S. muelleri*, and a 7,353 nt-long *ori* and a 4,498 nt-long *ter* fragment from *C. steedae.* Each purified fragment was then used as template to PCR-amplify dsDNA polynucleotide probes (referred to as Af/Sm/Cs-*ori*, and as Af/Sm/Cs*-ter*, respectively) (see Supplementary Table 2 and 3 for primers and PCR conditions).

All polynucleotide probes were chemically labeled with the Alexa Fluor 594 using the Ulysis Nucleic Acid Labeling Kit (ThermoFisher) following the same modifications to the manufacturers’ protocol as in (Barrero-Canosa et al., 2016).

### DNA fluorescence in situ hybridization (FISH)

Single *Robbea hypermnestrae* nematodes were rehydrated in PBS and nematode symbionts *Ca.* T. hypermnestrae were detached by sonication. Exponentially growing oral cavity symbionts were fixed in 3% or 4% PFA for 12-14 h at 4°C or for 1 h at room temperature, washed with PBS and stored in 1:1 ethanol/PBS. For the hybridization procedure, we followed a slightly modified version of the direct-gene FISH protocol ^60^. After letting the cell suspension dry onto a well of a Poly-L-lysine coated Epoxy-slide, cells were dehydrated in a series of increasing ethanol concentrations and permeabilized with freshly prepared lysozyme solution for 1 h on ice or at 37°C. Probes were diluted in hybridization buffer containing 35 or 45% formamide to final concentrations ranging 62-246 pg/ml and each probe was applied to the cells individually. Slides were transferred into a hybridization chamber and incubated for 40 min at 85°C and subsequently at 46°C for 2 h. Washing buffer was applied to the cells once briefly and once for 15 min at 48°C and, finally, cells were incubated in PBS and PBS supplemented with 5 µg/ml DNA stain Hoechst 33342 for 10 min at room temperature. Upon a quick wash in ddH_2_O and, subsequently, in 100% ethanol, cells were air-dried and mounted in 4.5 µl Vectashield mounting medium (Vector Labs) per microscopic slide well.

### DNA staining

Methanol-fixed nematodes or oral-cavity symbionts (fixed in 3% or 4% PFA for 12-14 h at 4°C or for 1 h at room temperature) were rehydrated and washed in PBS containing 0.1% Tween 20 (PBT), followed by incubation in 5 µg/ml Hoechst 33342 PBT for 15 min. After two washing steps to remove unbound DNA stain, worms were sonicated for 45 s to dissociate *Ca*. T. hypermnestrae from its host prior mounting. 1.5 ml of the bacterial suspension was mixed with 0.75 ml of Vectashield mounting medium (Vector Labs) and applied to a 1% agarose covered microscopy slide.

### Fluorescence microscopy

Slides containing symbiont cells subjected to DNA FISH were imaged using a Nikon Eclipse NI-U microscope equipped with a MFCool camera (Jenoptik). Images were acquired using the ProgRes Capture Pro 2.8.8 software (Jenoptik). Microscopic images were processed using the public domain program ImageJ ^61^ in combination with plugin ObjectJ ^62^ and Fil-Tracer ^32^ or MicrobeJ 5.13J ^63^. Cell outlines were traced and morphometric measurements recorded. Positions of the fluorescence foci (i.e., points of maximal fluorescent emissions) within the cell boundaries were measured and plotted as fraction of the normalized cell width and length of the cell that contained them. Automatic cell recognition was double-checked manually. For representative images, the background subtraction function of ImageJ was used, and brightness and contrast were adjusted for better visibility. Data analysis was performed using Excel 2021 (Microsoft Corporation), density plots were created with the ggplot2 ^64^ in R v.3.6.1 ^65^ and figures were compiled using Illustrator 2021 (Adobe Systems).

### Western blotting

The western blot against ParB was done as previously mentioned in Weber et al., 2019, with slight amendments. The primary antibody used was a custom peptide rabbit polyclonal anti-*A. filiformis* ParB antibody (Eurogentec) with a 1:1000 dilution. The secondary antibody used was a horseradish peroxidase-conjugated anti-rabbit secondary antibody (Amersham Biosciences) with a 1:10,000 dilution.

### Immunostaining

Deep-frozen methanol-fixed nematodes were rehydrated and washed in PBS containing 0.1% Tween 20 (PBT), followed by permeabilization of the bacterial peptidoglycan by a 15 min incubation with 0.1% (wt/vol) lysozyme at room temperature. Blocking was carried out for 1 hr in PBT containing 2% (wt/vol) bovine serum albumin (blocking solution) at room temperature. *Ca*. T. hypermnestrae was incubated with a 1:500 dilution of peptide rabbit polyclonal anti-*Ca*. T. oneisti ParB antibody (Eurogentec). All primary antibodies were incubated in blocking solution overnight at 4°C. Upon incubation with primary antibody (or without, in the case of the negative control) samples were washed three times in PBT and incubated with a 1:500 dilution of secondary Alexa 488-conjugated anti-rabbit antibody (Jackson ImmunoResearch, USA) in blocking solution for 1 hr at room temperature. Unbound secondary antibody was removed by three washing steps in PBT and thereupon incubated in 5 mg/ml Hoechst 33342 PBT for 15 min. After two washing steps to remove unbound DNA stain, worms were sonicated for 45 s to dissociate *Ca*. T. hypermnestrae from its host prior mounting. 1.5 ml of the bacterial suspension was mixed with 0.75 ml of Vectashield mounting medium (Vector Labs) and applied to a 1% agarose covered microscopy slide.

### Genome sequencing and assembly

For whole-genome sequencing, cells were lysed using lysozyme and Proteinase K, DNA extracted using phenol-chloroform and precipitated in two volumes of 100% ice-cold ethanol, 2.5 M ammonium acetate and 5 µg/ml glycogen. RNA was degraded by incubation with 10 mg/ml RNase A for 30 min at 37°C, and DNA cleaned up using DNA Clean & Concentrator-5 (Zymo). The library for Oxford Nanopore Technologies (ONT) sequencing was prepared using the ONT 1D ligation sequencing kit (SQK-LSK110) and sequenced on an R9.4 flow cell (FLO-MIN106) on a MinION for 48 h. Base calling was performed locally with ONT’s Guppy Basecalling Software guppy 5.0.11+2b6dbffa5 (dna_r9.4.1_450bps_sup.cfg), and resulting fastq-files were trimmed using using qcat (https://github.com/nanoporetech/qcat). For short read sequencing, sequencing libraries were prepared using the DNA NEB Ultra II FS (NEB) and sequenced on an Illumina NovaSeq SP with 100 bp single-end reads at the VBC NGS facility. The ONT data was assembled using canu v2.1.1 with genomeSize=2.5m ^66^, and polished with the long reads using 5 rounds of racon v1.4.3 (-m 8 -x −6 -g −8 -w 500, ^67^, reads mapped with minimap2 v2.7 -ax map-ont (Li, 2018) and samtools v1.9 ^68^, and one round of medaka v1.74.0 using the ‘consensus’ (--model r941_min_sup_g507) and ‘stitch’ modules (https://github.com/nanoporetech/medaka). The resulting assemblies were polished with the short reads using 5 rounds of pilon v1.22 ^69^ with bwa v0.7.16a mem ^70^ mapped reads. The genome completeness was assessed using CheckM version 1.0.18 ^71–73^ with the gammaproteobacterial marker gene set using the taxonomy workflow. Assembled contigs were annotated using Prokka v1.14.6 ^74^. Cluster of orthologous genes (COGs) were annotated using eggnog mapper v2.1.10 ^75^ against eggNOG 5.0 ^76^. Additional annotations were obtained through the MicroScope platform ^77^. For the ParB phylogeny, the alignment was performed with mafft v7.427 (L-INS-I mode) ^78^, and secondary structures predicted using ALi2D ^79,80^ and ESPript 3 ^81^ using the crystal structure of *Caulobacter crescentus* ParB (PDB# 6T1F). The maximum likelihood phylogeny was reconstructed using IQ-TREE v2.1.2 ^82^ with the best-fit model automatically selected by ModelFinder Plus ^83^ and 1,000 ultrafast bootstraps ^84^. The phylogeny was visualized in FigTree v1.4.4 (http://tree.bio.ed.ac.uk/software/figtree/).

### Prediction of the origin and of the terminus of DNA replication

The *ori* and *ter* positions were calculated based on GC skew and using Genskew (https://genskew.csb.univie.ac.at/) and smoothed using the lowess function (smootherspan = 0.005) of gplots v3.1.1 ^85^ in R 3.6.1 ^65^. *ParS* sequences were predicted using the conserved consensus *parS* (5’-NGTTNCANGTGNAACN-3’) ^58^ and *dif* sites using the betaproteobacterial consensus *dif* sequence (5’-HNNNBNNAYVAYNNDBVTTATGTHAANT-3’, ^59^. Sequencing coverage was obtained by mapping the raw genomic reads obtained for genome sequencing against the reference genomes using bowtie2 v2.4.4 ^86^ and samtools v1.9 ^68^. Mapped reads were counted and binned in 5 kb non-overlapping windows using bamCoverage of (standalone) deeptools v3.5.1 ^87^, and the resulting bigwig file converted to human-readable format using the bigWigToWig of UCSC v391. The coverage data was square transformed to better depict differences between *ori* and *ter*, and smoothed using the lowess function (smootherspan = 0.05). For illustration purposes (circular plots and 3C-seq heatmaps), the assemblies were reoriented to have the GC-skew-predicted *ori* in the middle of the genome sequence. Circular plots were generated using circos v0.69-8 ^88^ with Perl 5.028003 ^89^.

### Marker frequency analysis (MFA)

Cells were grown until an OD_600_ 0.6 (late exponential phase) and 1 (stationary phase) and collected by centrifugation at 16,000 g for 5 min at room temperature. Cells were lysed using lysozyme and Proteinase K, DNA extracted using Phenol-Chloroform and precipitated using ammonium acetate (2.5M final concentration), two volumes of 100% ice-cold Ethanol, and 5 µg/ml glycogen. RNA was degraded by incubation with 10 mg/ml RNase A for 30 min at 37°C, and DNA cleaned up using DNA Clean & Concentrator-5 (Zymo).

Sequencing libraries were prepared at the VBC NGS facility, and 150bp paired-end sequenced on an Illumina NovaSeq 6000 machine. Reads were adapter-trimmed and filtered using trimmomatic v0.39 (ILLUMINACLIP:adapters.fa:2:30:10 LEADING:3 TRAILING:3 MINLEN:80 SLIDINGWINDOW:4:15 ^90^ and prinseq-lite v0.20.4 (-min_qual_mean 30 -min_len 25) ^91^. PhiX contaminant sequences were removed using bbduk.sh of bbmap v38.90 (k=31 hdist=1) ^92^, as well as further contaminants (human, animal, plant, fungi) removed against repeat-masked references (minid=0.95 maxindel=3 bwr=0.16 bw=12 quickmatch fast minhits=2 qtrim=rl trimq=10 untrim). Reads were mapped against the reference chromosomes generated in this study using bowtie2 v2.4.4 ^86^, and index-sorted using samtools v1.9 ^68^.

Reads were counted using the bamCoverage program of (standalone) deeptools v3.5.1 ^87^ in non-overlapping 1kb windows and the resulting bigwig file converted to human-readable format using the bigWigToWig of UCSC v391. The count data was smoothed using the lowess function (smootherspan = 0.1) and plotted using ggplot2 ^64^ in R v.3.6.1 ^65^, and the ratio of the GC-skew predicted *ori* to *ter* coverage (number of non-normalized mapped reads) calculated.

### Chromosome conformation capture (3C-seq)

3C libraries were prepared as previously described ^7,19^. For growth on solid medium, *C. steedae*, *A. filiformis* and *S. muelleri* were plated from glycerol stock onto BSTSY agar plates and grown at 37°C for 24 h and 48 h, after which they were scraped off and transferred to BSTSY/FBS containing 6% f.c formaldehyde, fully resuspended, and fixed at room temperature at 120 rpm, followed by 30 min more at 4°C. Glycine (250 mM f.c.) was added, and the cells incubated 30 min at 4°C at 120 rpm. After fixation, cells were collected by centrifugation for 10 min at 3,500 g at 4°C, resuspended in fresh medium, collected again by centrifugation, and resuspended in 1 ml fresh medium before final collection by centrifugation. Cell pellets were stored at −70°C. For liquid cultures, *A. filiformis* was grown until OD_600_ of 0.3 (exponential phase) or 1 (stationary phase). For blocking transcription or translation, cells grown overnight on plate were incubated with Rifampicin (25 µg/ml f.c.) or Chloramphenicol (25 µg/ml) for 30 min prior to cell fixation and collection.

For digestion, cell pellets were thawed on ice for 10 min, resuspended in 500 µl Tris 10 mM EDTA 0.5 mM (TE) (pH 8), and lysed using 4 µl Ready-lyse lysozyme (Epicentre) for 20 min at 37°C, 300 rpm. SDS (0.5% f.c.) was added and incubated for 10 min at room temperature. 500 µl lysed cells were then transferred to a tube containing 4.5 ml digestion buffer (1% Triton X-100, 1x CutSmart buffer (New England Biolabs)), and 100 µl cells (non-digested control) were transferred to a tube containing 0.9 ml of digestion buffer. 100U of HpaII (New England Biolabs) were added to the 5 ml digestion mix. Both tubes were incubated for 3 h at 37°C. After digestion, 1 ml of digestion mix was put on ice (digested control) together with the non-digested control. The remaining digestion mix was split into 4 parts, centrifuged for 20 min at 20,000 g, and pellets were carefully resuspended in 1 ml water each, and pooled into 7 ml ligation buffer (1X ligation buffer 3 (New England Biolabs; without ATP), 1 mM ATP, 1 mg/ml BSA, 500 U of T4 DNA ligase 5 U/µl (ThermoFisher)). Ligation was carried out at 16°C for 4 h, followed by incubation together with the two control tubes overnight at 65°C with proteinase K (0.25 mg/ml f.c.) and EDTA (6 mM f.c.). The next day, DNA of the ligated sample was precipitated with 1/10 volume of 3 M sodium acetate (pH 5.2) and two volumes of isopropanol, and after one hour at −80°C, DNA was pelleted by centrifugation for 30 min at 10,000 g at 4°C and resuspended in 500 µl 1X TE buffer. DNA in all tubes (ligation and two controls) were extracted once with 400 µl phenol-chloroform pH 8.0 (vortexed for 1 min and centrifuged for 5 min 16,000 g at 4°C), precipitated (see above; 30 min at −80°C, followed by centrifugation for 20 min at 16,000 x g at 4°C), washed with 400 µl 80% cold ethanol, and diluted in 30 µl 1X TE buffer supplemented with RNase A (1 mg/ml f.c.; ThermoFisher). Tubes containing ligated DNA were pooled. A 1% agarose gel was run with aliquots of the ligated and two control DNA to check for efficiency of digest and ligation.

Ligated DNA (5 µg) was adjusted with water to 130 µl, and sheared using a Covaris S220 (Duty cycle 5, Intensity 5, cycles/burst 200, time 60 s for 4 cycles), and the DNA purified using the QIAquick PCR purification kit (40 µl elution buffer; Qiagen), and DNA end-repaired with 1X T4 DNA ligase buffer (New England Biolabs), 0.4 mM dNTP mix, 15 U T4 polynucleotide kinase (New England Biolabs), 5 U Klenow DNA polymerase (Roche), and incubated for 30 min at RT. DNA was purified using MinElute columns (Qiagen). A-tailing was done with 1X NEBuffer 2 (New England Biolabs), 0.2 mM dATP, and 15 U Klenow exo (New England Biolabs), for 30 min at 37°C before inactivation for 20 min at 65°C, and DNA again purified using MinElute columns (Qiagen). Sequencing adapters were ligated with 0.5 mM custom-made adapters, 2.10^6^ U T4 DNA ligase, and 1X T4 DNA ligase buffer, for 2 hours at RT, followed by inactivation for 20 min at 65°C. DNA fragments were purified in a size from 400 to 900 bp using a PippinPrep apparatus (SAGE Science). Each library was amplified in 12 cycles with 1X Phusion Buffer (ThermoFisher), 0.2 mM dNTPs, 0.2 µM Illumina primers PE1.0 (5’-AATGATACGGCGACCACCGAGATCTACACTCTTTCCCTACACGA-3’) and PE2.0 (5’-CAAGCAGAAGACGGCATACGAGAT-3’), 3 µl of 3C library, and 1U of Taq Phusion polymerase (ThermoFisher; 98 °C 30 s, 12 cycles of 65°C 30 s, and 72°C 30 s, and 72°C for 7 min). The PCR reaction was purified using MinElute PCR purification kit (Qiagen), and libraries sequenced on an Illumina NextSeq500 with 75 bp paired-end reads.

### Generation of contact maps

Raw contact maps were built as described previously^18^. Briefly, each read was assigned to a restriction fragment. Non-informative events such as self-circularized restriction fragments, or uncut co-linear restriction fragments were discarded. The chromosome of each bacterium was divided into 5 kb bins and the frequencies of contacts between genomic loci for each bin were assigned. Contact frequencies were visualized as heatmaps. Raw contact maps used for comparative analyses were built using 1 million valid reads. Raw contact maps were normalized using Iterative correction and eigenvector decomposition (ICE) ^93^To facilitate visualization, normalized contact matrices are visualized as log matrices. A Gaussian filter of H=0.5 is used when plotting the matrices. To quantify the frequency of interactions in the secondary diagonals (Supplementary Figure 6), we applied to our data a method based on the filtering of significant 3C contacts and developed as described^55^.

### Identification of boundaries in contact maps

To identify boundaries we applied the Frontier Index (FI) method as described in detail in ^42^ and summarized in ^19^. Briefly, the FI method allows to identify loci associated with demarcations, or frontiers, in the 3C contact maps that can occur at any scale, meaning that the method is useful to identify domain organization irrespective of the underlying scale. Specifically, at a given locus (let say *i*), two frontiers can be found at most, each one corresponding to a side of the locus – one frontier indicates that the neighboring loci on the same side of the locus *i* tend to make more contacts with each other with respect to neighboring loci separated by the same genomic distances but located on each side of the locus. Visually, identified frontiers can be vertical (or upstream) or horizontal (downstream) (see e.g. Fig. 4) ^42^.

### Loop analysis

To facilitate analysis, normalized bins interacting with less than 10% of the total number of bins were considered empty due to multi-mapping reads such as multiple rRNA operons and removed from the analysis. This corresponded to approximately 10% of the total number of bins in S. muelleri. In practice, we removed the corresponding row and columns of the 3C-seq matrices and analyzed the resulting smaller matrices.

Loop detection was performed as following. We first slightly smoothed 3C-seq matrices using a Gaussian filter with width 0.5. Next, we computed a 4C profile for each bin. Peaks in these profiles were good candidates for loops such that we extracted the highest peaks of each profile, where the height of a peak was defined as the difference of 4C value between the peak and the maximum value of the two flanking minima. The resulting histogram of the heights could be divided into an exponential distribution for small values and a strong deviation from this exponential distribution for large values. In this context, we considered a conservative threshold value to distinguish the two sets of values. 4C peaks higher than this threshold were then considered as loop. Eventually, looping regions were defined as maximal sets of contiguous bins such that any two consecutive bins were involved in at least one loop with a.

### Transcriptomics

All organisms were grown in the same conditions as those applied for the 3C-seq experiments in three biological replicates each as for the 3C-seq experiments. 500 µl liquid culture were mixed well with 1 ml RNAprotect Bacteria Reagent (Qiagen) and incubated for 5 min, and cells grown on solid medium scraped off and resuspended in 1 ml thereof. Cells were collected by centrifugation for 10 min at 5,000 g, and cell pellets stored at −70°C until the next day.

Total RNA was extracted using the RNeasy Mini Kit (Qiagen) for low cell numbers, including the on-column DNA digestion. Eluted RNA (30 µl) was treated with 2 U of DNase for 30 min using the TURBO DNA-free kit (Invitrogen). RNA was cleaned up using the RNA Clean & Concentrator-25 kit (Zymo) including the on-column DNA digestion. Absence of DNA was checked via PCR (35 cycles) against a ∼200 bp fragment of *gyrB*. RNA was quality checked at the VBC NGS facility (all samples had a RIN > 9), rRNA depleted using an in-facility developed protocol, and reverse-stranded sequencing libraries were prepared using the NEBNext Ultra II Directional RNA Library Prep Kit (New England Biolabs). Libraries were sequenced on an Illumina NovaSeq 6000 machine, using 100 bp single-end reads (*A. filiformis*) and 150 bp paired-end reads (*S. muelleri, C. steedae*). Reads were adapter-trimmed and filtered using trimmomatic v0.39 (ILLUMINACLIP:adapters.fa:2:30:10 LEADING:3 TRAILING:3 MINLEN:80 SLIDINGWINDOW:4:15; (Bolger et al., 2014)) and prinseq-lite v0.20.4 (-min_qual_mean 30 -min_len 25) ^92^, as well as further contaminants (human, animal, plant, fungi) removed against repeat-masked references (minid=0.95 maxindel=3 bwr=0.16 bw=12 quickmatch fast minhits=2 qtrim=rl trimq=10 untrim). Reads mapping to rRNA were removed using sortmerna v4.1.0 ^94^ against the pre-built sortmerna databases and the organism-specific rRNA sequences. Reads were mapped against the reference genome generated in this study using bowtie2 v2.4.4 ^86^, Q>30 filtered and index-sorted using samtools v1.9 ^68^. Reads were counted using featureCounts of the subread-package v2.0.0 ^95^ in single-end mode (-s 2) or in paired-end mode (-s 2 -p -B -C) against the “gene” features of the reference genomes’ gff-files.

To show transcription levels along the chromosome, trimmed reads were mapped against the reference chromosomes using bowtie2 v2.4.4 ^86^, and index-sorted using samtools v1.9 ^68^. Per-basepair coverage was determined using samtools depth (-a -d 0) ^68^, and binned in 5 kb bins using using a custom script in R v3.6.1 ^65^. Counts were normalized to the sum of counts per sample and square-root transformed. All plots were made using matplotlib v3.5.1^96^ in python v3.7.12 ^97^.

For differential expression analysis, raw per-gene counts and metadata (i.e., conditions) were imported using DESeqDataSetFromMatrix in DESeq2 package v1.30.1 ^98^ in R v4.0.3 ^65^, and differential expression analysis carried out using the function ‘DESeq’, and results extracted using the function ‘results’ with padj.cutoff = 0.05 and lfc.cutoff = log2(2). For each sample, raw per-gene counts were divided by gene length, and TPM values calculated by multiplying each normalized count value by 10^6^, divided by the sum of all counts per sample. Per-sample TPM values were averaged per condition. Genes were classified into expression level categories by considering the distribution parameters as indicated in Supplementary Fig. 10.

