## Supplementary information for "Stable chromosome configuration and loop-based polarization in animal symbionts" (from another version of this preprint: version 1, 2023-12-23)

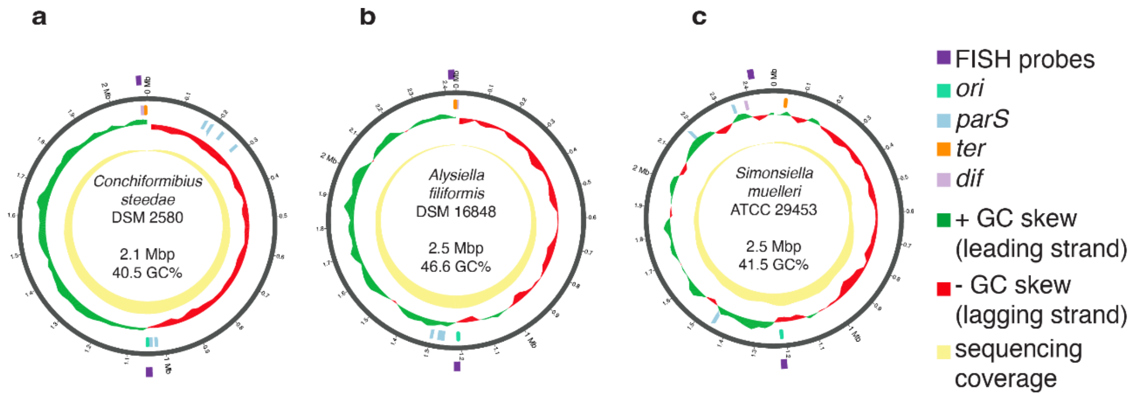


**Supplementary Figure 1. Circular chromosome maps of *Conchiformibius steedae* DSM 2580 (a), *Alysiella filiformis* DSM 16848 (b) and *Simonsiella muelleri* ATCC 29453 (c)**. Inner circle: position in megabases (Mb, black); middle circle: GC skew (green for positive, red for negative); inner circle: sequencing coverage (yellow). Colored highlights represent the position of the *ori* (light green), *parS* sequences (light blue), *ter* (orange), *dif*-site (light purple), and DNA FISH probes (dark purple) targeting the predicted *ori* and *ter*, respectively.

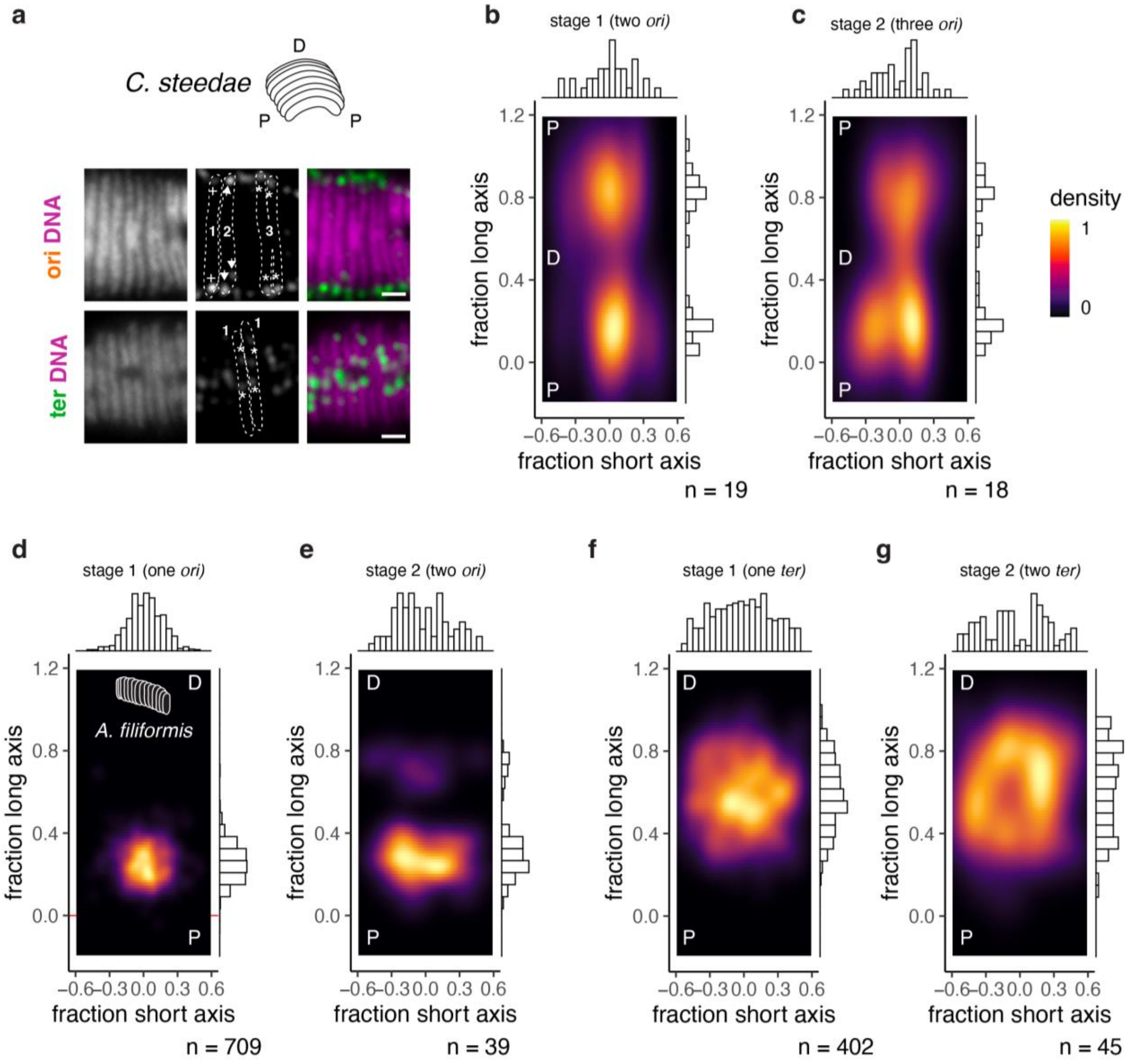


**Supplementary Figure 2. Lateral segregation of *C. steedae* and *A. filiformis*.** (a) Images of representative *C. steedae* filaments subjected to DNA FISH with an *ori*-specific probe (upper panels) and with a *ter*-specific probe (bottom panels). Cells with two foci, three foci or four foci were arbitrarily referred to as stage 1, 2 and 3, respectively. Pluses, arrowheads or asterisks indicate foci in stage 1, 2 or 3 cells, respectively. White dotted cell outline is the cell shape deduced from phase contrast images (leftmost panels). Overlays are displayed in the rightmost panels. Scale bars are 1 µm. (b) DNA FISH-based subcellular localization of *C. steedae ori* foci in 19 stage 1 cells, (c) of *C. steedae ter* foci in 18 stage 3 cells, (d) of *A. filiformis ori* in 709 stage 1 cells, (e) of *A. filiformis ori* in 39 stage 2 cells, (f) of *A. filiformis ter* in 402 stage 1 cells, (g) of *A. filiformis ter* in 45 stage 2 cells.

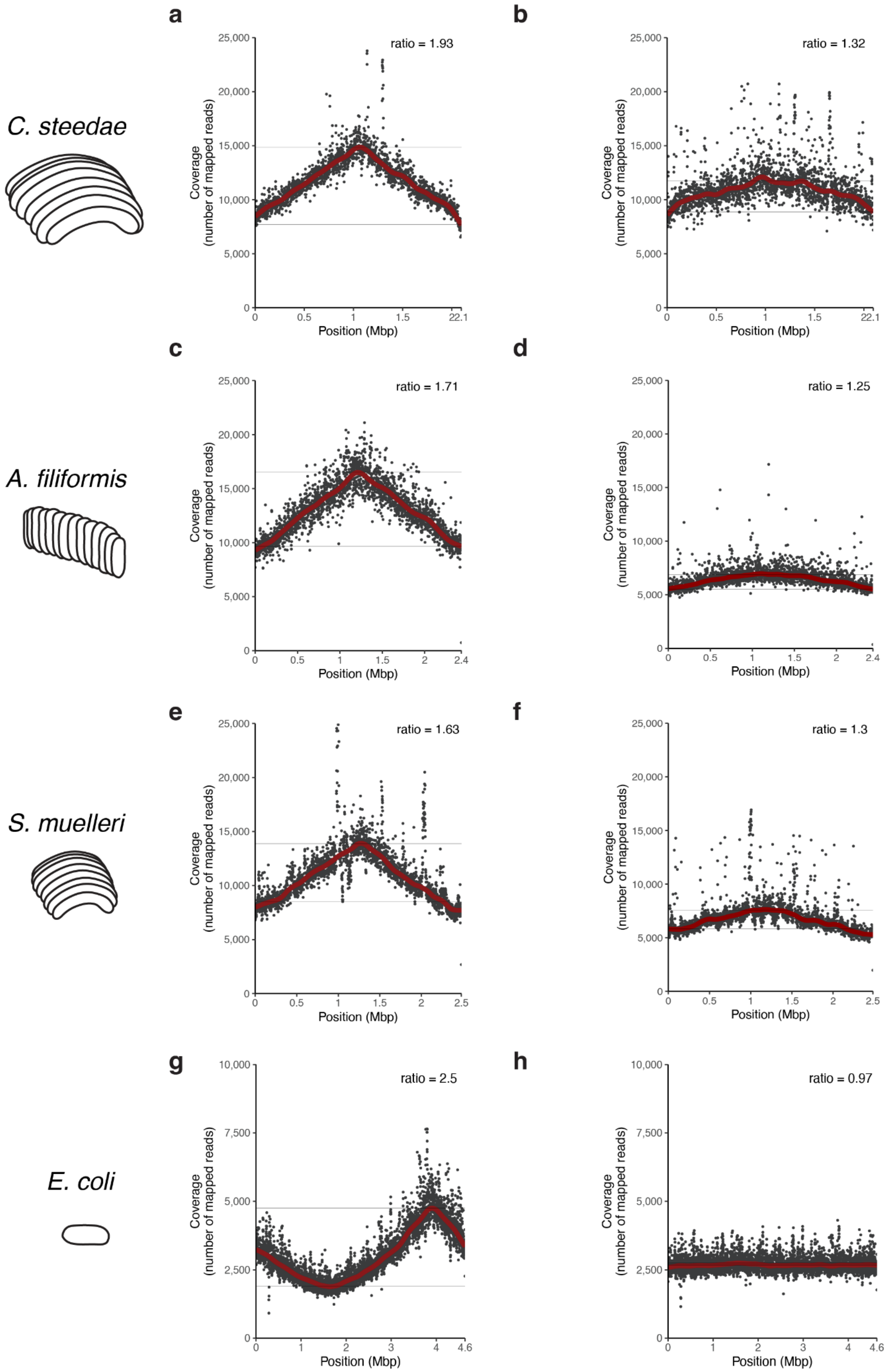

a
b
c
d
e
f
g
h
C. steedae
A. filiformis
S. muelleri
E. coli
ratio = 1.93
ratio = 1.32
ratio = 1.71
ratio = 1.25
ratio = 1.63
ratio = 1.3
ratio = 2.5
ratio = 0.97
Coverage (number of mapped reads)
Position (Mbp)

**Supplementary Figure 3. Marker frequency analysis (MFA) of (a) *C. steedae* in exponential phase and (b) stationary phase, (c) *A. filiformis* in exponential phase and (d) stationary phase, (e) *S. muelleri* in exponential phase and (f) stationary phase, and (g) *E. coli* in exponential phase and (h) stationary phase.** The x axis represents the genomic coordinates of the reference genome, and the y axis the coverage expressed in number of mapped reads. The coverage along the chromosome (black dots) is smoothed (red line), and horizontal lines represent the coverage at the *ori* (highest coverage) and the *ter* (lowest coverage), which was used to calculate the *ori*-*ter* ratio. For comparison, we performed MFA of *Escherichia coli* in (g) exponential phase and (h) stationary phase based on published data (Ivanova et al., 2015).

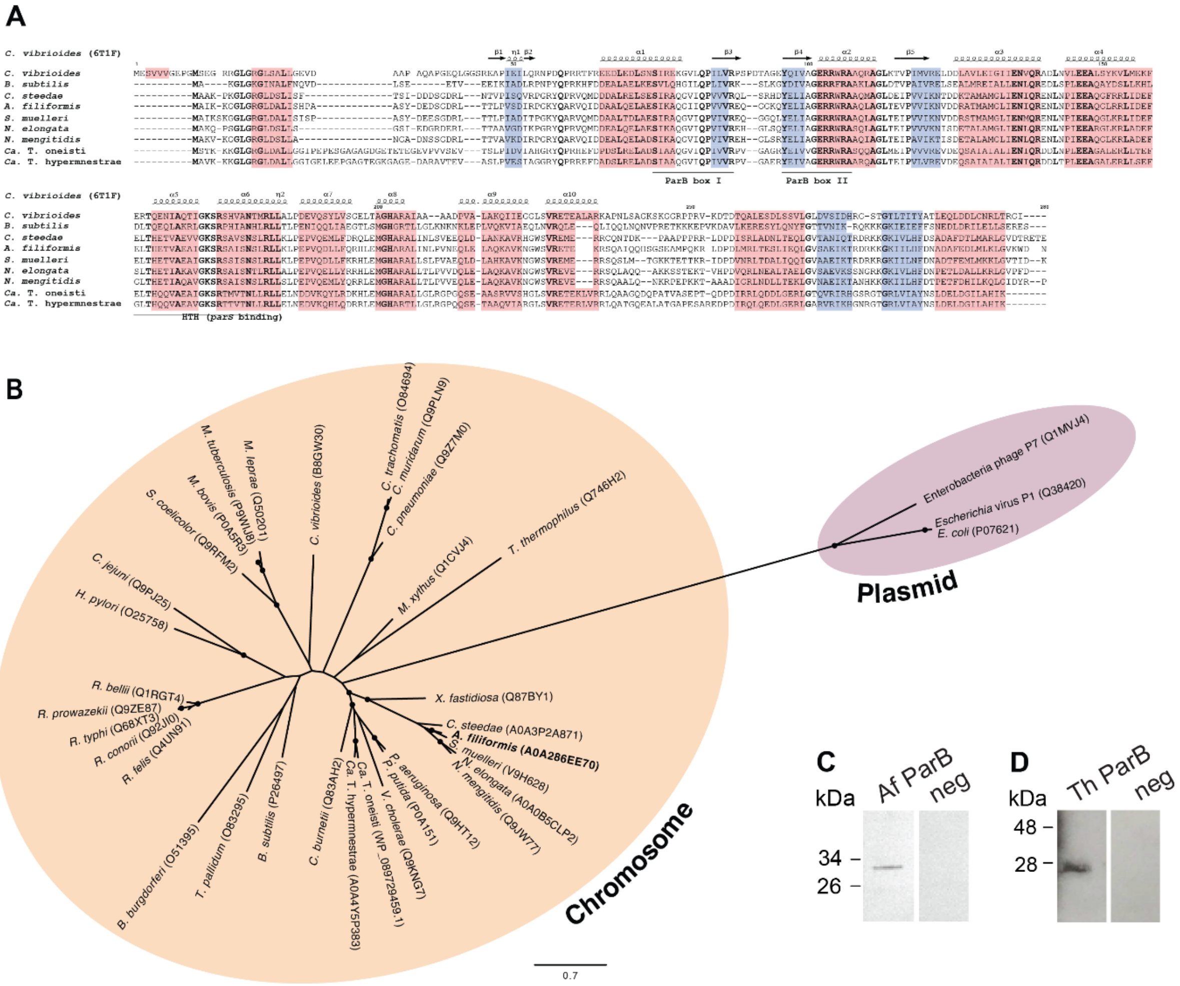


**Supplementary Figure 4. Structural alignment of ParB, phylogenetic placement and protein expression in *A. filiformis* and *Ca.* T. hypermnestrae.**

(A) Secondary structure prediction of representative bacterial ParB. Red indicates predicted alpha helices whereas blue indicates predicted beta strands. The experimental secondary structure elements are inferred from the *Caulobacter vibrioides* ParB and depicted above each sequence block. Prediction was performed using PSIPRED and Ali2D. (B) Unrooted phylogenetic tree including ParB of nematode and oral cavity symbionts, as well as diverse other bacterial ParB. Sequences underlayed in orange facilitate chromosome segregation and in purple plasmid segregation. The analysis is based on a MAFFT alignment of full-length proteins (data not shown) and estimated under the Q.pfam+G4 model using ML analysis (IQ-TREE) with node support calculated by UltraFast bootstraps. Scale bar represents 0.7% estimated sequence divergence. Closed circles represent values ≥95% based on 10,000 replicates. (C) Western blots of protein extracts of *A. filiformis* and *Ca.* T. hypermnestrae probed with a specific anti-ParB antibody with the secondary antibody only (neg). Numbers indicate apparent MWs expressed in kDa.

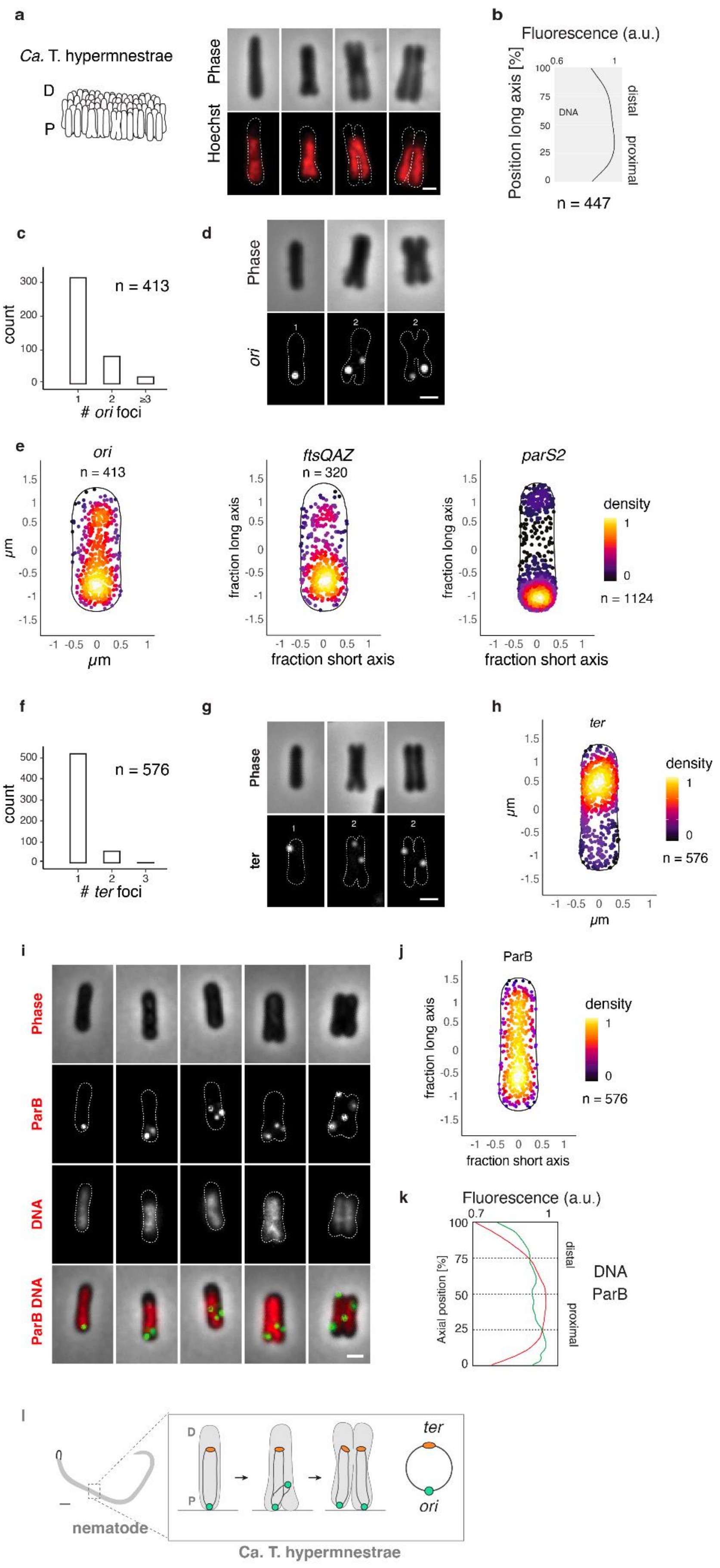

a
Ca. T. hypermnestrae
D
P
Phase
Hoechst
b
Fluorescence (a.u.)
Position long axis [%]
DNA
distal
proximal
n = 447
c
count
n = 413
### ori foci
d
Phase
ori
e
ori
n = 413
ftsQAZ
n = 320
parS2
density
n = 1124
μm
fraction long axis
fraction short axis
f
count
n = 576
### ter foci
g
Phase
ter
h
ter
density
n = 576
μm
i
Phase
ParB
DNA
ParB DNA
j
ParB
density
n = 576
fraction long axis
fraction short axis
k
Fluorescence (a.u.)
Axial position [%]
distal
proximal
DNA
ParB
l
D
P
ter
ori
nematode
Ca. T. hypermnestrae

**Supplementary Figure 5. DNA localization, chromosome configuration and ParB localization in *Ca.* T. hypermnestrae.** (a) Hoechst 33342 staining of fixed *Ca.* T. hypermnestrae cells. Cells are arranged from the youngest to the oldest from left to right. Upper panels display phase contrast images of cells and lower panels their corresponding DNA fluorescence (white dotted line represent cell outlines). (b) Averaged DNA fluorescence (a.u.) plotted against the cell length (right). (c-f) Histograms showing number of (c) *ori* foci (n = 413) and (f) *ter* foci (n = 576) per cell. (d, g) Confocal microscope images of representative filaments of *A. filiformis* subjected to (d) *ori* DNA FISH or (g) *ter* DNA FISH; phase contrast images are displayed in upper panels and corresponding FISH (white) fluorescence images in bottom panels. (e) Subcellular localization of 413 *ori*, 320 fts*QAZ* and 1,124 *pars2* foci (from left to right) and (h) subcellular localization of 576 *ter* foci. (i) Representative images showing five *Ca.* T. hypermnestrae cells arranged from the youngest to the oldest from left to right. Panels display phase contrast images, ParB and DNA fluorescence and corresponding overlays (from top to bottom). White dotted lines represent cell outlines. (j) Subcellular localization of symbiont ParB foci of 576 cells immunostained with an anti-ParB antibody. Scale bars are 1 µm. (k) Averaged ParB and DNA fluorescence (a.u.) plotted against the cell length (n = 82). (l) Model of *ori*-*ter* chromosome configuration and lateral chromosome segregation in the monoploid *Ca*. T. hypermnestrae*.* Scale bar is 1 µm in (a,d,g, i) and 0.1 mm in (l).

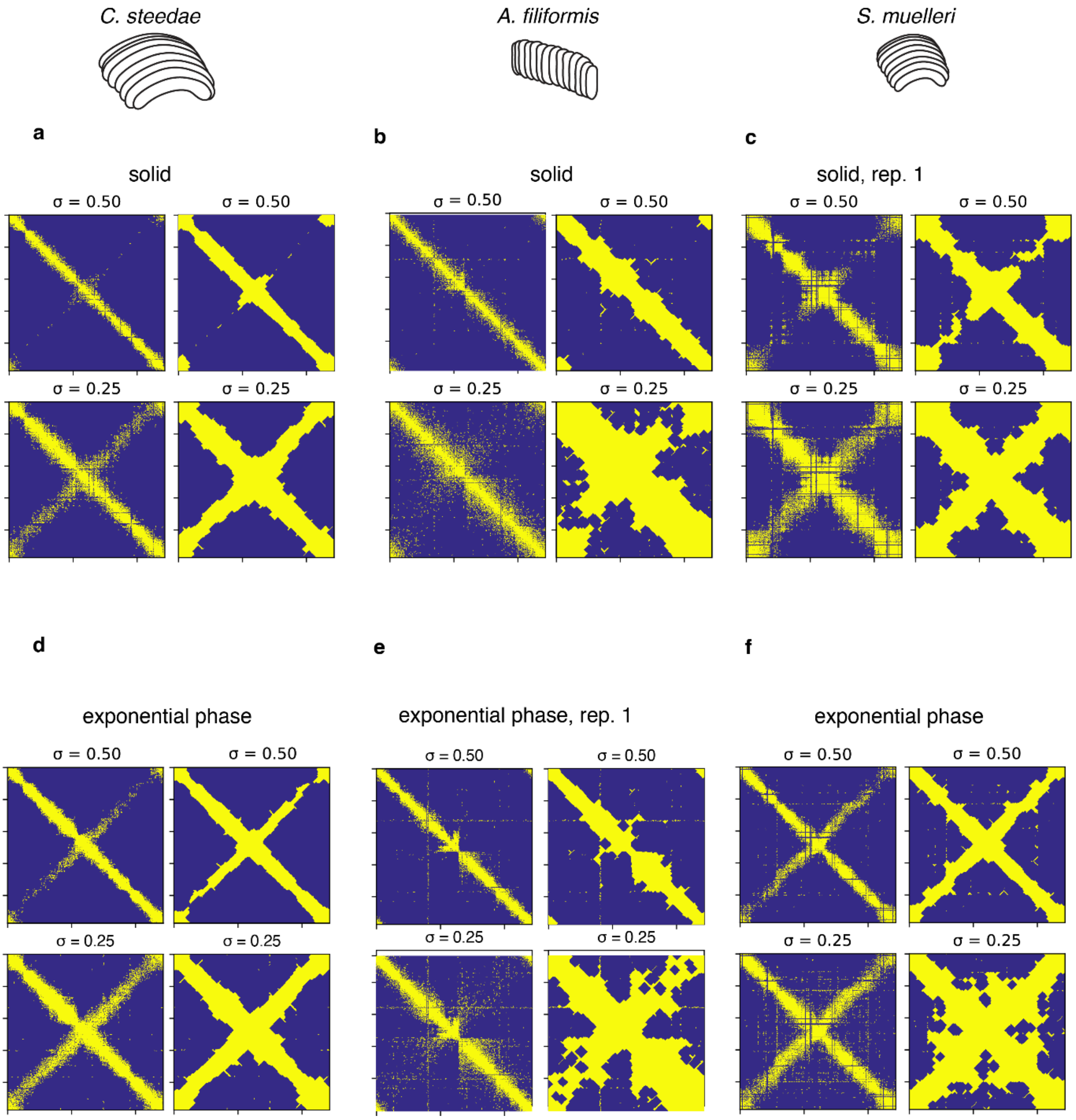


**Supplementary Figure 6. Contact map analysis of the primary and secondary diagonal in *C. steedae*, *A. filiformis* and *S. muelleri*.** Contact maps of *C. steedae* (a and d), *A. filiformis* (b and e) and *S. muelleri* (c and f) were generated after setting the threshold for significant interactions at 0.5x, and 0.25x the standard deviation above the median to the non-cleaned, normalized contact maps obtained in each condition. Contact frequencies above or below this threshold were assigned a value of 1 or 0, respectively, generating a binary contact map in which significant interactions are plotted in yellow and non-significant interactions in blue (left row of each panel). Additionally, by connecting the signal in the binary maps by connecting 5 elements considered as significant using a diamond shape of 15 to fill out empty points comprised by the connected elements (right row of each panel).

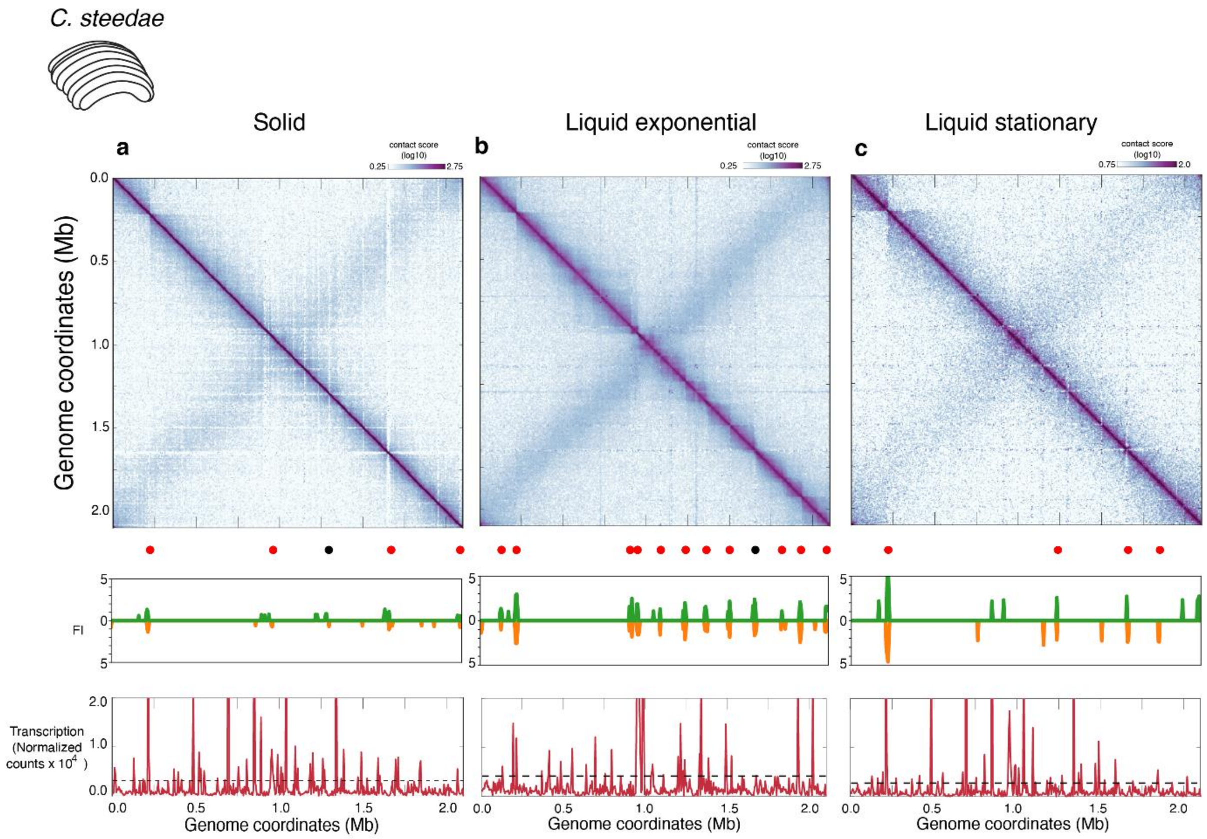


**Supplementary Figure 7. Normalized chromosome contact maps, frontier index (FI) analyses, boundaries and transcriptomic correlation in *C. steedae*.** Contact maps of *C. steedae* grown (a) on solid medium, (b) in liquid medium, exponential phase or (c) in liquid medium, stationary phase. Below each map, the track shows CID borders, position of highly expressed genes (HEGs) and rRNA operons if their presence correlates with a CID border. Below each map, the black or red circles show the position of highly expressed genes (HEGs) and rRNA operons if their presence correlates with a CID border. Below each track, the Frontier Index (FI) shows frontiers with upstream (green) or downstream (orange) bias of interaction and the normalized RNA-Seq counts in 5 kb bins show the transcriptomic data obtained in the same conditions that the contact map. The dashed line represents the threshold used to identify highly expressed region.

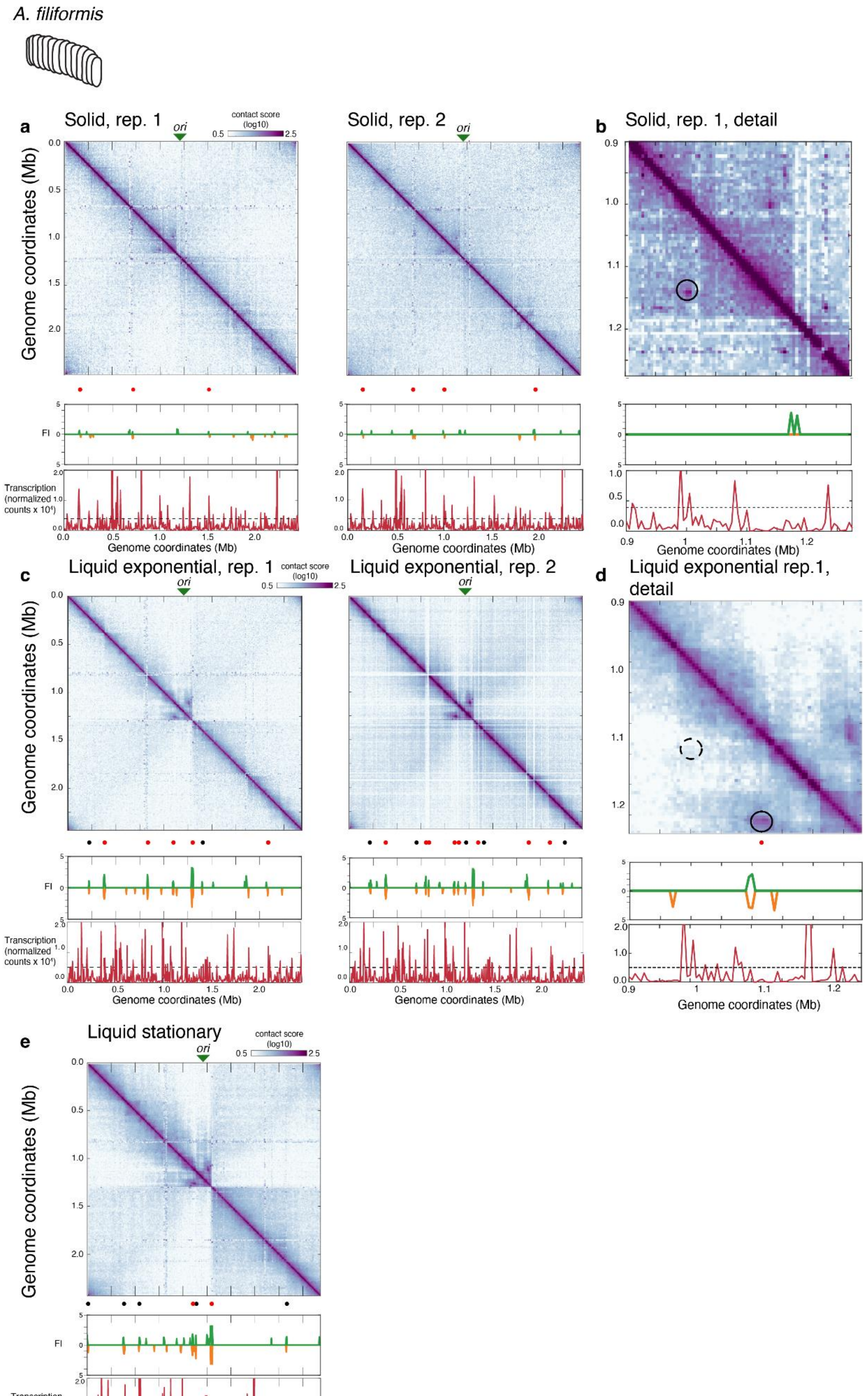

A. filiformis
a
Solid, rep. 1
Solid, rep. 2
ori
contact score (log10)
0.5
2.5
Genome coordinates (Mb)
FI
Transcription (normalized counts x 10⁴)
b
Solid, rep. 1, detail
c
Liquid exponential, rep. 1
Liquid exponential, rep. 2
d
Liquid exponential rep.1, detail
e
Liquid stationary

**Supplementary Figure 8. Normalized chromosome contact maps, frontier index (FI) analyses, boundaries and transcriptomic correlation in *A. filiformis*.** (a) Contact maps of *A. filiformis* grown on solid medium replicate 1 (left panel) and replicate 2 (right panel). (b) Detail (0.9-1.28 Mb) of (a) showing the loop situated on the right of the *ori*. (c) Contact maps of *A. filiformis* grown in liquid medium until exponential phase (replicate 1, left panel and replicate 2, right panel). (d) Detail (0.9 and 1.28 Mb) of (c). The black circle indicates the loop and the dashed circle indicates the position of the loop shown in (b). (e) Contact map of *A. filiformis* grown in liquid medium until stationary phase. Below each map, the track shows CID borders, position of highly expressed genes (HEGs) and rRNA operons if their presence correlates with a CID border. Below each map, the black or red circles show the position of highly expressed genes (HEGs) and rRNA operons if their presence correlates with a CID border. Below each track, the Frontier Index (FI) shows frontiers with upstream (green) or downstream (orange) bias of interaction and the normalized RNA-Seq counts in 5 kb bins show the transcriptomic data obtained in the same conditions that the contact map. In all panels, dashed line in the transcription plots represents the threshold used to identify highly expressed regions.

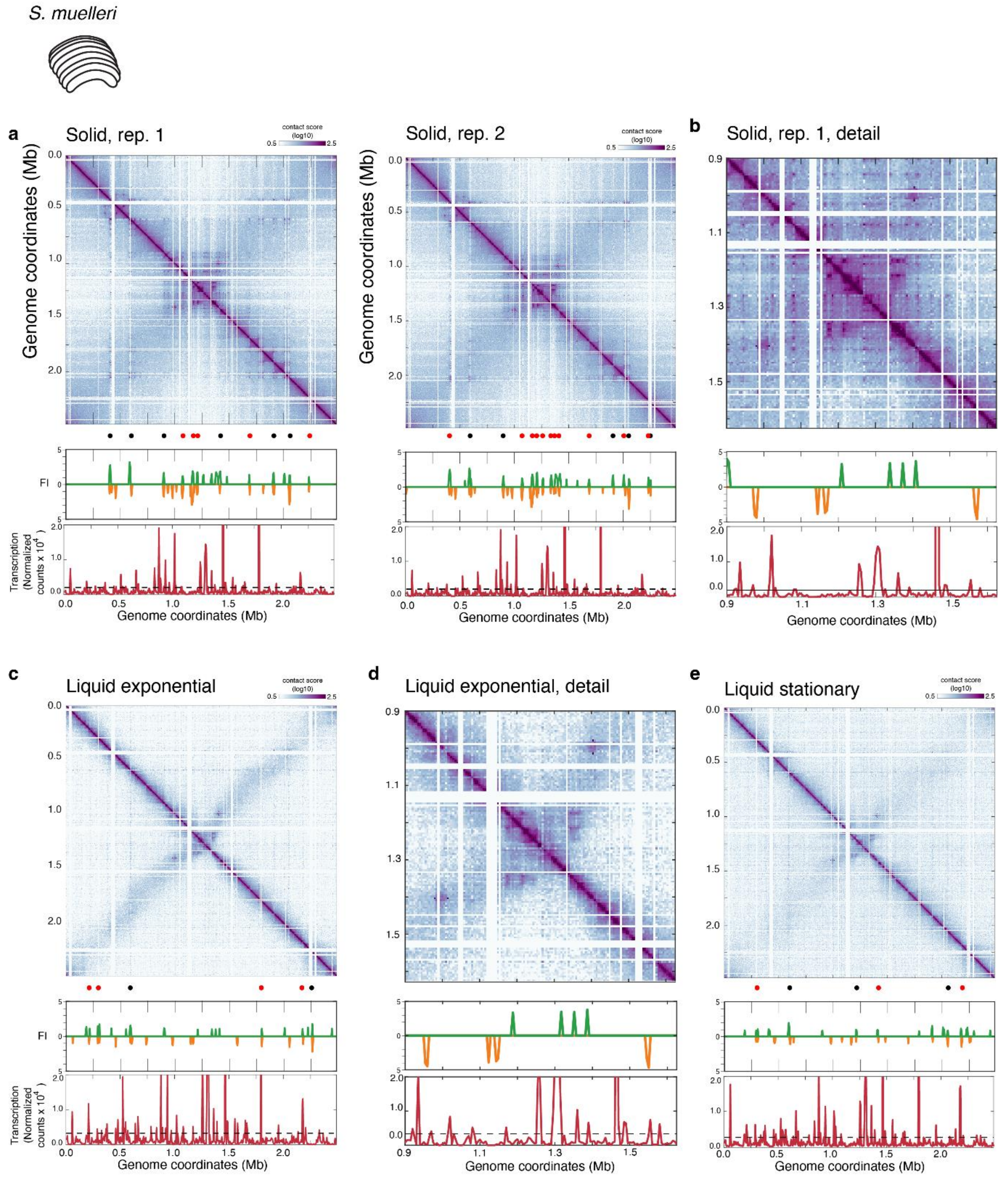


**Supplementary Figure 9. Normalized chromosome contact maps, frontier index (FI) analyses, boundaries and transcriptomic correlation in *S. muelleri*.** (a) Contact map of *S. muelleri* grown on solid medium (replicate 1, left panel) and replicate 2 (right panel), (b) Detail (0.9-1.625 Mb) of the contact map shown in (a). (c) Contact map of *S. muelleri* grown in liquid medium until exponential phase. (d) Detail (0.9-1.625 Mb) of the contact map shown in (c). (e) Contact map of *S. muelleri* grown in liquid medium until stationary phase. . Below each map, the track shows CID borders, position of highly expressed genes (HEGs) and rRNA operons if their presence correlates with a CID border. Below each map, the black or red circles show the position of highly expressed genes (HEGs) and rRNA operons if their presence correlates with a CID border. Below each track, the Frontier Index (FI) shows frontiers with upstream (green) or downstream (orange) bias of interaction and the normalized RNA-Seq counts in 5 kb bins show the transcriptomic data obtained in the same conditions that the contact map. In all panels, dashed lines in the transcription plots represent the threshold use to identify highly expressed regions.

a

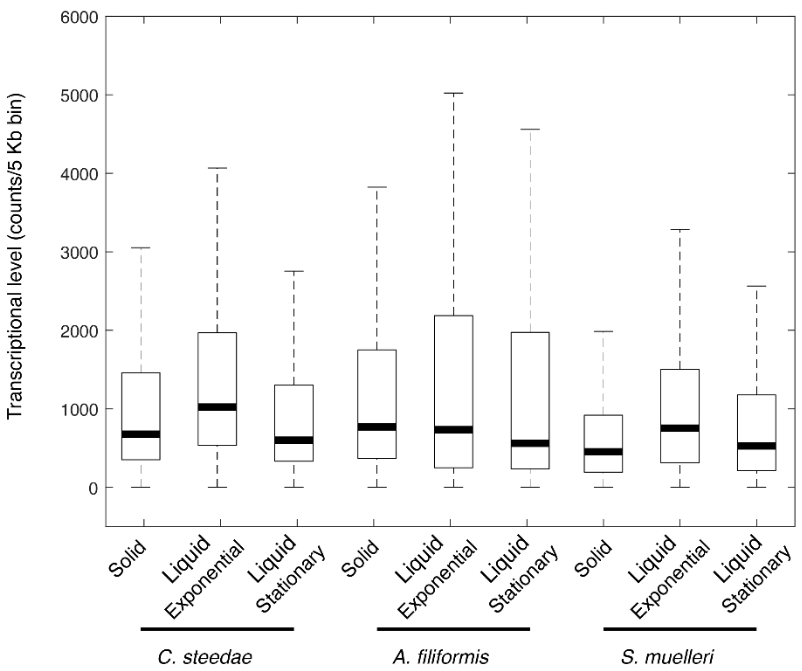


b

Categories of transcription level:

| | | |
|---|---|---|
| Cat_4 | Upper limit | Very high |
| Cat_3 | Between Q3 & upper limit | High |
| Cat_2 | Between median & Q3 | Medium |
| Cat_1 | Between Q1 & median | Low |
| Cat_0 | Between 0 and Q1 | No/poor |

c

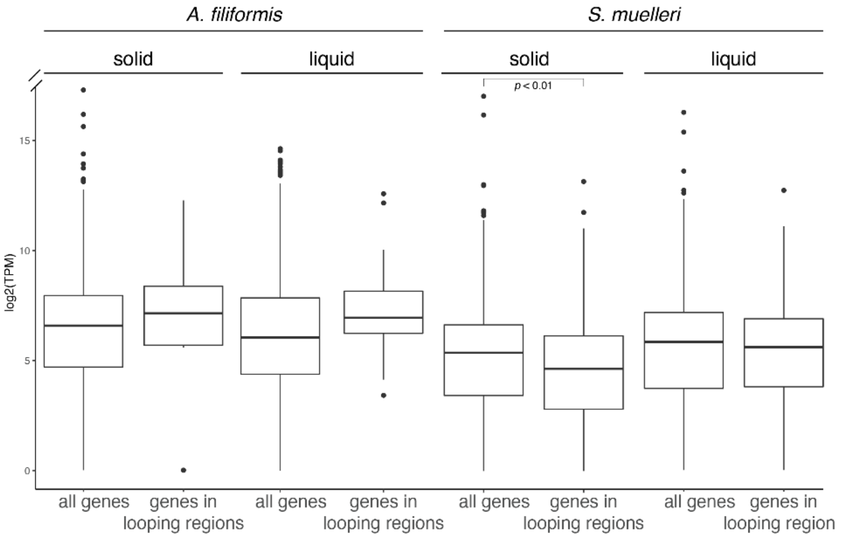


**Supplementary Figure 10. Transcriptomic data of oral cavity symbionts**. (a) For *C. steedae*, *A. filiformis* and *S. muelleri* transcriptomes were binned at 5 kb resolution and represented as boxplots. The boxplots represent the first quartile (Q1), median and third quartile (Q3). The upper whisker extends from the hinge to the largest value no further than 1.5 * the inter-quartile range (IQR, i.e., distance between the first and third quartiles) from the hinge. The lower whisker extends from the hinge to the smallest value at most 1.5 * IQR of the hinge. For simplicity, outliers are not represented but considered for the analyses. (b) Categories used to classify the expression level of each bin on the chromosome. (c) Boxplots of expression levels (as $\log_2$(TPM)) of all genes or genes positioned in looping regions, for *A. filiformis* and *S. muelleri*, on solid medium and in liquid. For better readability, the y-axis has been truncated.

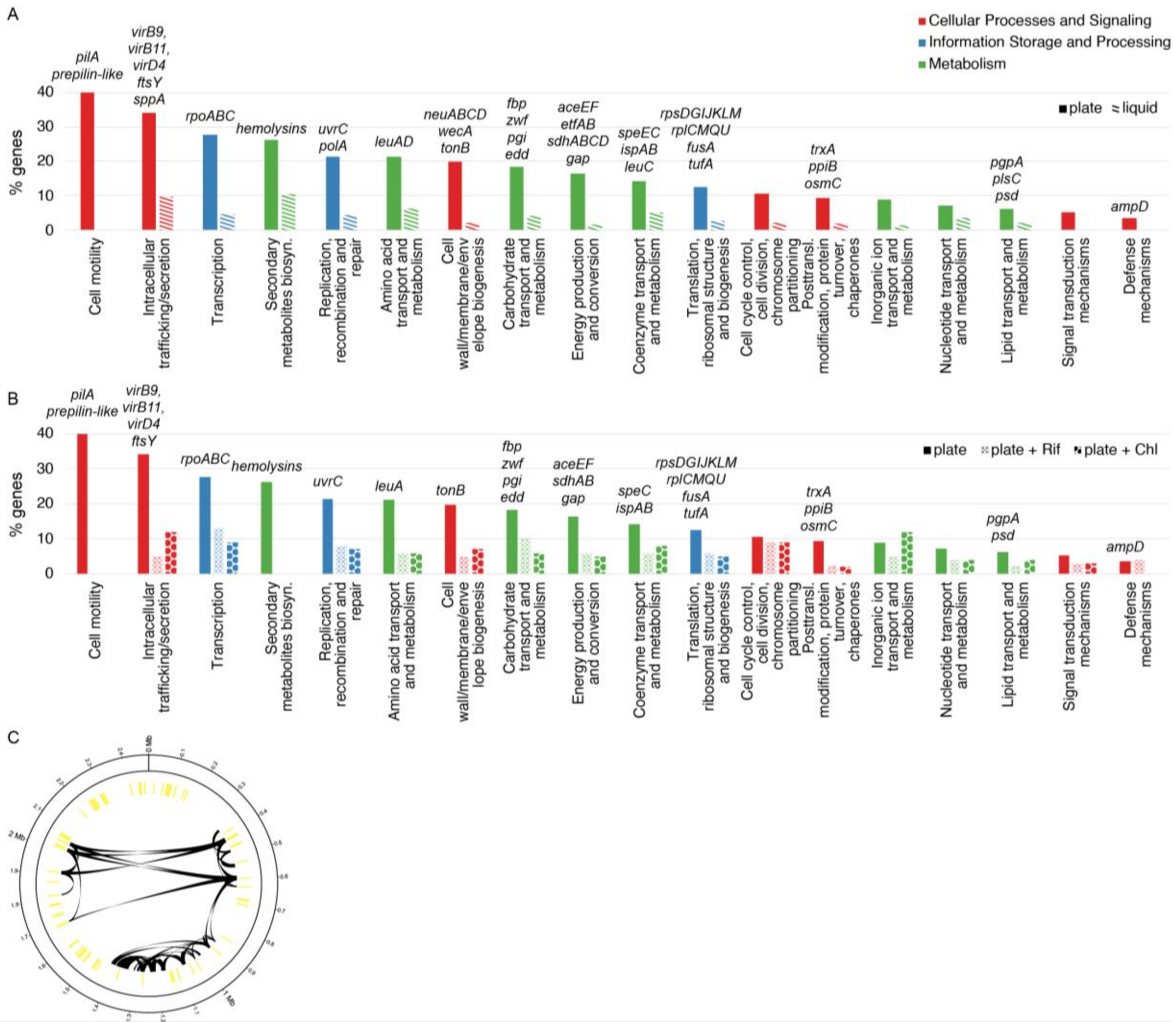


**Supplementary Figure 11. EggNOG-based annotation of genes situated in looping regions in *S. muelleri* grown on solid medium or in liquid medium and circular representation of the *S. muelleri* chromosome with looping regions and position of identified insertion sequences.** (A) Histogram of *S. muelleri* COG categories of genes in looping regions on solid medium (plate) and liquid medium (liquid) represented as % of the total number of genes in each category. (B) Histogram of *S. muelleri* COG categories of genes in looping regions on solid medium (plate), solid medium with 25 µg/ml Rifampicin (plate +Rif) and in solid medium with 25 µg/ml Chloramphenicol (plate +Chl) represented as % of the total number of genes in each category. Solid-specific COG categories are indicated in bold. Name abbreviations of genes discussed in the main text are listed besides the COG categories they belonged to. The colors of the bars indicate the three COG categories ‘Cellular Processes and Signaling’ (red), ‘Information Storage and Processing’ (blue), and ‘Metabolism’ (green). See Supplementary Data Set 1 for additional information, e.g., gene names, predicted intracellular localization, etc. (C) Insertion sequences (yellow dashes) were predicted using ISEScan [99].

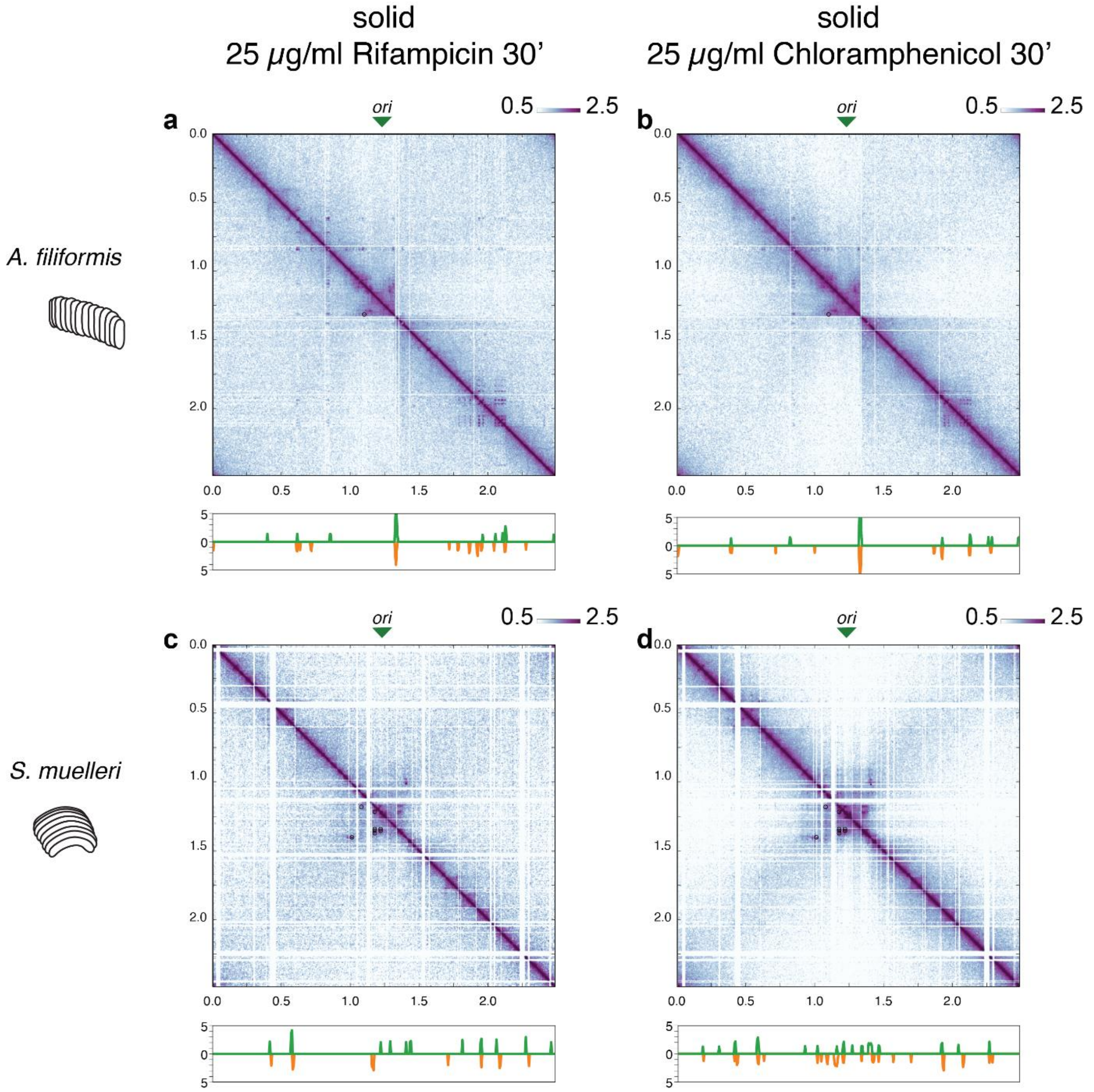


**Supplementary Figure 12. Normalized chromosome contact maps of *A. filiformis* and *S. muelleri* grown on solid medium with 25 µg/ml Rifampicin or 25 µg/ml Chloramphenicol for 30 min*.*** (a) Contact map of *A. filiformis* grown on solid medium with 25 µg/ml Rifampicin and (b) with 25 µg/ml Chloramphenicol. (c) Contact map of *S. muelleri* grown on solid medium with 25 µg/ml Rifampicin and (b) with 25 µg/ml Chloramphenicol. Below each panel is the frontier index analysis.

**Table S1.** DNA FISH foci counts.

| Species | *ori* number per cell (%) | | | | *ter* number per cell (%) | | | |
|---|---|---|---|---|---|---|---|---|
| | **1** | **2** | **3** | **>4** | **1** | **2** | **3** | **>4** |
| Th ($n_{ori}$ =413; $n_{ter}$ =576) | 76 | 20 | 5 | - | 90 | 10 | - | - |
| Af ($n_{ori}$ =748; $n_{ter}$ =447) | 98 | 2 | - | - | 96 | 4 | - | - |
| Sm ($n_{ori}$ =188; $n_{ter}$ =293) | 29 | 47 | 20 | 4 | 61 | 30 | 7 | 2 |
| Cs ($n_{ori}$ =215; $n_{ter}$ =236) | 27 | 44 | 21 | 7 | 25 | 32 | 22 | 22 |

**Table S2.** DNA FISH primers.

| Target | Probe No. | Primer | Sequence (5'-3') | Product (nt) |
|---|---|---|---|---|
| ***A. filiformis ori region*** (9,232 nt) | 1 | Af_Ori1F | GGTTTTCAATGCCGTATCT | 300 |
| | | Af_Ori1R | GACAATTTGTCCAAAGATGAGG | |
| | 2 | Af_Ori2F | CGCAGCCAATTCAATATGAC | 300 |
| | | Af_Ori2R | CAACAAGCAGGTTCAGAAG | |
| | 3 | Af_Ori3F | GTGGTGCAGTTTCAGATAGT | 300 |
| | | Af_Ori3R | AAACTCTACAACGGTAGCG | |
| | 4 | Af_Ori4F | TGTCAAACCAATCATGACG | 300 |
| | | Af_Ori4R | AAGACACGTCTTTGCGTAT | |
| | 5 | Af_Ori5F | GCAATCACATAGCGACCTT | 300 |
| | | Af_Ori5R | AACCAATCCCTTGTCTGAA | |
| | 6 | Af_Ori6F | TTTAAGCCAGCCTTTTGGAT | 300 |
| | | Af_Ori6R | GGATTTTGAGTTTGACCCG | |
| | 7 | Af_Ori7F | GCTTCAGCTTGAGATACTG | 300 |
| | | Af_Ori7R | AGACATTTTGGAAGCGGTAA | |
| | 8 | Af_Ori8F | TGGAATAGAAGCCAACTCAG | 300 |
| | | Af_Ori8R | TATCGCGGTATCAGTGTTG | |
| | 9 | Af_Ori9F | ACGCTAGATACATCTACGC | 300 |
| | | Af_Ori9R | GATTTTGGGTCCTCGTGG | |
| | 10 | Af_Ori10F | ACACATCTACTGATTCGTCA | 300 |
| | | Af_Ori10R | CAGTTCCAATTCTTTGACCAAC | |

| | | | | |
|---|---|---|---|---|
| ***A. filiformis ter region*** (6,778 nt) | 1 | Af_Ter1F | TTGGTTTACCCCATTAGCG | 300 |
| | | Af_Ter1R | CATGTCAAGGCAGCTTTG | |
| | 2 | Af_Ter2F | ATGTGGCAGCAATTTCAG | 300 |
| | | Af_Ter2R | GAACACGAATTGCCCTTTTA | |
| | 3 | Af_Ter3F | ACGCAAATACGCCAAAAAC | 300 |
| | | Af_Ter3R | TACATTTGGGGCGACAAA | |
| | 4 | Af_Ter4F | AACTTTGCACCACATATTCG | 300 |
| | | Af_Ter4R | GCCGTTTTGTTCAAGTGT | |
| | 5 | Af_Ter5F | TTATAATGCGCCCAACAATG | 300 |
| | | Af_Ter5R | GCACTTCGCATTAAAAACAC | |
| | 6 | Af_Ter6F | TGACGACAAAGTGGAACG | 300 |
| | | Af_Ter6R | AACAACCACCAAGCCAAT | |
| | 7 | Af_Ter7F | CCTGAAACAAAATCCGCC | 300 |
| | | Af_Ter7R | ATTCATGCGGATTTTGACC | |
| | 8 | Af_Ter8F | TGGAAGAGTCCACCAGTT | 300 |
| | | Af_Ter8R | AAAACAACCACATTGGCG | |
| | 9 | Af_Ter9F | GCAACAACACGCCATTTT | 300 |
| | | Af_Ter9R | TTCCCCTGCCCCATTATTA | |
| | 10 | Af_Ter10F | CGCACAGCAATTCAAAATCT | 300 |
| | | Af_Ter10R | GTGTGAATGTGATACAAGCG | |

| | | | | |
|---|---|---|---|---|
| ***S. muelleri ori* region** (8,158 nt) | 1 | Sm_ori_1F | CAATGTAGATGCCGCTGAA | 300 |
| | | Sm_ori_1R | CAACGATAGGTCACATTTGC | |
| | 2 | Sm_ori_2F | GATTTAGCAGCAGAACAACC | 300 |
| | | Sm_ori_2R | GCCACAAATATCCACTACGT | |
| | 3 | Sm_ori_3F | CTATATATTCATTCGCTGGACG | 300 |
| | | Sm_ori_3R | TTACCGCTTTTACGTGGG | |
| | 4 | Sm_ori_4F | ATTGATTACGTTGGCTCATG | 300 |
| | | Sm_ori_4R | TTTTACGAGCAAGCTGTCT | |
| | 5 | Sm_ori_5F | GTGCTGATTTACGACGAAG | 300 |
| | | Sm_ori_5R | CACCAATTGAGAAGTACGC | |
| | 6 | Sm_ori_6F | CCAAACCCACTACACATAAT | 300 |
| | | Sm_ori_6R | GTTTTGGTGATCCTGAAAC | |
| | 7 | Sm_ori_7F | CAAAATTAGCCAGCGGAATC | 300 |
| | | Sm_ori_7R | TTTTGGGGCAGATTGATTC | |
| | 8 | Sm_ori_8F | TATTACCTAACTTTCGCTCCG | 300 |
| | | Sm_ori_8R | AAAAAGGAGTGTGTACCGA | |
| | 9 | Sm_ori_9F | TACTATCCTGCATAC CTGCA | 325 |
| | | Sm_ori_9R | AATTTGGTGGCTACGAATG | |
| | 10 | Sm_ori_10F | CATTTGTGCAGTTTGGTTAG | 300 |
| | | Sm_ori_10R | GTGAAAATGTCCATGAATCC | |

| | | | | |
|---|---|---|---|---|
| ***S. muelleri ter* region** (8,187 nt) | 1 | Sm_ter_1F | GATTTGCGGTCAATCGTT | 300 |
| | | Sm_ter_1R | GGGCAAATTCAATTCAGACC | |
| | 2 | Sm_ter_2F | ATGAAACCATCTGAACCGT | 300 |
| | | Sm_ter_2R | CATTGATGCACTGGTTATTG | |
| | 3 | Sm_ter_3F | GATGATTTCACGGATTTTAGCC | 300 |
| | | Sm_ter_3R | CGTTCTCAATTTGTAGACCAAG | |
| | 4 | Sm_ter_4F | TCGTTAATTCAGGCGTGTC | 300 |
| | | Sm_ter_4R | GGACGGTTGGACTTATTCG | |
| | 5 | Sm_ter_5F | TTGTCCGTCTAAATAAGGTGTC | 300 |
| | | Sm_ter_5R | TTGGACAAACTCAAACATGG | |
| | 6 | Sm_ter_6F | CACCGAATGATCAGGAATG | 300 |
| | | Sm_ter_6R | GGTTTAAAAGCGAGAGACG | |
| | 7 | Sm_ter_7F | TTCCTTGTGGGTCTAATACTTG | 300 |
| | | Sm_ter_7R | CAGAGCCATTAAGTAAATCAC G | |
| | 8 | Sm_ter_8F | AGCATCAACACGTACGAAA | 300 |
| | | Sm_ter_8R | GATTATCGGCATGAGACGG | |
| | 9 | Sm_ter_9F | GCGTAGGATTATTCTCATTAGC | 314 |
| | | Sm_ori_10R | GTGAAAATGTCCATGAATCC | |

| | | | | |
|---|---|---|---|---|
| **Ca. T. hypermnestrae *ori* region** (2,858 nt) | 1 | Th_dnaA_start | TCCGTGGAAATCTGGAATCAATGT | 306 |
| | | Th_ori_1R | TGTGGATTTTTCTTGAGGTGTGG | |
| | 2 | Th_ori_2F | CCGCGAGCAACTTAATTACCG | 344 |
| | | Th_ori_2R | GCCTGCGAAGAACTGAATATCA | |
| | 3 | Th_ori_3F | TCGAATCCAAACAGCAAATCGT | 315 |
| | | Th_ori_3R | GTACGTTTCTGGCAAATTCGAG | |
| | 4 | Th_ori_4F | GGACATGCTTGCTGCTCAG | 315 |
| | | Th_ori_4R | CATTATGCTGTCAGGGTTCTGAAC | |
| | 5 | Th_ori_5F | TTCAGTGCCAGCAATTTCGCC | 310 |
| | | Th_ori_5R | CCGGATAAAGTCGTCCTTTGCC | |
| | 6 | Th_ori_6F | CGGAACCGATCTAGAAATAGAGG | 304 |
| | | Th_ori_6R | CGCCATCGAAAATGCAGTCTT | |
| | 7 | Th_ori_7F | CAGGACGTTCGCTATTATCT | 334 |
| | | Th_ori_7R | CATAGGCAACTTGCTCCGGA | |
| | 8 | Th_ori_8F | AGATACACTGAAGCAGGCATTG | 317 |
| | | Th_ori_8R | GTCTCATCGGCATTACTACAAAG | |
| | 9 | Th_ori_9F | GACACATGATTTTGAAGGTAGAG | 216 |
| | | Th_dnaA_stop | CAGCCTCTCAGTGTCACCGT | |

| | | | | |
|---|---|---|---|---|
| ***Ca. T. hypermnestrae ter* region** (3,239 nt) | 1 | Th_dif_F1 | GTAAGGGGATTGCCAGGCTTG | 311 |
| | | Th_dif_R1 | GTGGACGACACGGTGCATTT | |
| | 2 | Th_dif_F2 | CACGATGCCCAGCGTCATG | 333 |
| | | Th_dif_R2 | CTCTCGACCAAGGAGGTACTTGC | |
| | 3 | Th_dif_F3 | GTCGCCCGCTTGTCGATATCG | 314 |
| | | Th_dif_R3 | GAGAACCTGACCGGCGTTTATT | |
| | 4 | Th_dif_F4 | ATGAAATCGGTATCCGTACGG | 318 |
| | | Th_dif_R4 | GAATTTCTCCGAGGCGCCG | |
| | 5 | Th_dif_F5 | AGCGTCAGAAACCCGATGG | 323 |
| | | Th_dif_R5 | CTGCTGATCATTGTGGTCGCC | |
| | 6 | Th_dif_F6 | GCCAGCGGCAGGAGATGC | 324 |
| | | Th_dif_R6 | GGTTGTCGATCTGATTGAGGACCC | |
| | 7 | Th_dif_F7 | GTCAGCGCATCCCCCTCG | 319 |
| | | Th_dif_R7 | GGCTGGTGATTGCAGGGACC | |
| | 8 | Th_dif_F8 | AGCGATGGCCCAAAATCCAAG | 305 |
| | | Th_dif_R8 | ATGATGTCCCCACGGGATAATTCG | |
| | 9 | Th_dif_F9 | CATGCAGACCTTTGTCGCGC | 330 |
| | | Th_dif_R9 | GGCTTCCTCCTTGATGTGCTCG | |
| | 10 | Th_dif_F10 | GTGCTGGCGGCGATGGGA | 326 |
| | | Th_dif_R10 | GCCTGTGCTTAAGTGGCTGGT | |
| | 11 | Th_dif_F11 | ACCGCGATCACGCGCTGA | 325 |
| | | Th_dif_R11 | AAACCGGAACCTGCTGACCA | |
| | 12 | Th_dif_F1.2 | CGGATTACGCGGATGGGGTAA | |
| | | Th_dif_R11.2 | TATTAGCAGATTGCGACGCGCG | |

| | | | | |
|---|---|---|---|---|
| ***Ca. T. hypermnestrae parS2* region** (3,524 nt) | 1 | Th_parS2_1f | CTCTACCTGTTTGATCGCGTTCG | 319 |
| | | Th_parS2_1r | CAAGGTCTTCAGCGAGATGTACG | |
| | 2 | Th_parS2_2f | TCGGCGGATTCGAGCCACA | 290 |
| | | Th_parS2_2r | CTGAGTCGTCTCCTCTCCATGG | |
| | 3 | Th_parS2_3f | GAGCCTCCCAGCATCATTTCC | 313 |
| | | Th_parS2_3r | ATGCCGGCCATCAGGTCTTC | |
| | 4 | Th_parS2_4f | CCATCGGGTCCGTCCAGGA | 321 |
| | | Th_parS2_4r | CGCTTGTTTCGAGCCTCTATTCC | |
| | 5 | Th_parS2_5f | GTTATTTATACTACAGCGCCGAGCG | 330 |
| | | Th_parS2_5r | GCCAGGTTTTTCGCCACGG | |
| | 6 | Th_parS2_6f | CGAGGACAAAGCCAAGGTCG | 316 |
| | | Th_parS2_6r | GGTGAGGAAGGTGCCGATG | |
| | 7 | Th_parS2_7f | GGATGGCCGACAGCGGTAAG | 305 |
| | | Th_parS2_7r | AATGACGTGCTTGATGGTGCC | |
| | 8 | Th_parS2_8f | AACGACTCGATCGACGGCA | 313 |
| | | Th_parS2_8r | GATCTTAATCATGCGCGATTCTCC | |
| | 9 | Th_parS2_9f | GACGACCACAAGCTGGTACG | 320 |
| | | Th_parS2_9r | GCCTCGATCATCTCCTCGG | |
| | 10 | Th_parS2_10f | GATCCGCTCGGTCGTCAGTG | 311 |
| | | Th_parS2_10r | CCGCTCAGGCATCCTCCTC | |
| | 11 | Th_parS2_11f | GGATGTCCCGGATTGTGCGG | 333 |
| | | Th_parS2_11r | CTGAGCAGGACGTTATACCGG | |

| | | | | |
|---|---|---|---|---|
| **Ca. T. hypermnestrae *ftsQAZ* region** (3,690 nt) | 1 | Th_ftsQAZ_F1.2 | CTGCCCCTGATCCGCCTGGA | 335 |
| | | Th_ftsQAZ_R1.1 | AATGGTTCAAGGCTGGTCTCC | |
| | 2 | Th_ftsQAZ_F2.1 | CCGGACAAGCGATGAACAGG | 329 |
| | | Th_ftsQAZ_R2.1 | CAGGCGATCCGGCCAGTGA | |
| | 3 | Th_ftsQAZ_F3.1 | GAGCTCCGGGTGGTGGAAC | 323 |
| | | Th_ftsQAZ_R3.1 | GGGTAGACCCGGATAAACCTGGC | |
| | 4 | Th_ftsQAZ_F4.1 | CCGTCGGTGATCGATATGCG | 319 |
| | | Th_ftsQAZ_R4.1 | TCCTTCACCTCGCCCACCAG | |
| | 5 | Th_ftsQAZ_F5.1 | CGACGAGGGGATCGAGATTATCG | 312 |
| | | Th_ftsQAZ_R5.1 | TGGGGCAGGATGTGCAGGAT | |
| | 6 | Th_ftsQAZ_F6.1 | AATTCATCATCGACAATCAGGAAGGG | 308 |
| | | Th_ftsQAZ_R6.1 | GCCGGCGATGGGAATCACG | |
| | 7 | Th_ftsQAZ_F7.1 | GATCAGGTGACCAACGATATCGCC | 286 |
| | | Th_ftsQAZ_R7.1 | AGCTGCCGCCGGTCATGAC | |
| | 8 | Th_ftsQAZ_F8.1 | CCAAGATGGCGGGGCTCAT | 361 |
| | | Th_ftsQAZ_R8.1 | CCGTTTTGCTCGATGGTTATCTGATT | |
| | 9 | Th_ftsQAZ_F9.1 | ATGTTTGAACTCATGGATACCAACGG | 397 |
| | | Th_ftsQAZ_R9.1 | TCACTACCGCCACGGTCAGG | |
| | 10 | Th_ftsQAZ_F10.1 | CCAAGCCCTTTGCGTTCGAG | 349 |
| | | Th_ftsQAZ_R10.1 | AAGGGGCTGTTGATGGCCG | |
| | 11 | Th_ftsQAZ_F11.1 | GCTGGAGGATATCGACCTGGC | 336 |
| | | Th_ftsQAZ_R11.1 | CCCTTGTTGCGATAGACCGCC | |

| | | | | |
|---|---|---|---|---|
| ***C. steedae ori region*** (7,353 nt) | 1 | C_Ori1F | TCAATTTCAAAAACCGGCTC | 300 |
| | | C_Ori1R | GATAGAACACATTATGTCTGCG | |
| | 2 | C_Ori2F | GCATGAGTGTGAAAGTAGC | 300 |
| | | C_Ori2R | AAAAACCTGTGCATTCCAG | |
| | 3 | C_Ori3F | AACTGTGGTGTTCAACGTTA | 300 |
| | | C_Ori3R | AGTTTGCCTTTCAGTTCAG | |
| | 4 | C_Ori4F | GTAGCCACGGGTTCATTGA | 331 |
| | | C_Ori4R | ACCACTTAGCCTAATTATCGG | |
| | 5 | C_Ori5F | AGGTTAAACAGTCAGTGGTAG | 300 |
| | | C_Ori5R | TTTTTCGATCAAAGACGGG | |
| | 6 | C_Ori6F | TTTACCGAGCAATATGTCG | 300 |
| | | C_Ori6R | GCGTACAAAGCGGTTAATC | |
| | 7 | C_Ori7F | GTGGTGCTGATTTACGATGA | 300 |
| | | C_Ori7R | CAGACGTTGACTCAATGTG | |
| | 8 | C_Ori8F | GCGTCCCACGGATAATTAAT | 300 |
| | | C_Ori8R | CGAATATTACGCCAACGAAAG | |
| | 9 | C_Ori9F | GCTTACCATACCTTCAAGC | 300 |
| | | C_Ori9R | CGGCAAAAAGAAAGTTTGAG | |
| | 10 | C_Ori10F | GTTAGACATAAACCCGACATGA | 300 |
| | | C_Ori10R | CACCATGCTACTTGGTCTG | |
| | 11 | C_Ori11F | AGAAAGGAAACCCCATGTC | 300 |
| | | C_Ori11R | GATTAAAGGTGGTAAATTCGCG | |

| | | | | |
|---|---|---|---|---|
| ***C. steedae ter region*** (4,498 nt) | 1 | C_Ter1F | TTGCCCGACAACATTACTAC | 300 |
| | | C_ Ter1R | CTGTTTGCGGTTCATATAGG | |
| | 2 | C_Ter2F | GAAGGTTTTGAACGCATTCA | 300 |
| | | C_Ter2R | CGAACTTTACACCTTTCCTTC | |
| | 3 | C_Ter3F | CGTAGATTTAAACGCTGACCT | 300 |
| | | C_Ter3R | TAAGCCATTTCCACACTCG | |
| | 4 | C_Ter4F | GCCGACAGTATTTGCTTAC | 300 |
| | | C_Ter4R | CTTTGTGTTGCCCGATATAATC | |
| | 5 | C_Ter5F | CCAAAGAGCAACTGGGTAT | 300 |
| | | C_Ter5R | TGTCCAGTTGAAAATCTTCC | |
| | 6 | C_Ter6F | GCTTACCTTATTAGATGTGG | 300 |
| | | C_Ter6R | GTAGGTTTTGTAGGGCTTT | |
| | 7 | C_Ter7F | GGTGGCGTATTGCTTAACT | 300 |
| | | C_Ter7R | GTGTAAAGGTTCTAAATGGGA C | |
| | 8 | C_Ter8F | ACACCTTTTTATCGGTAGC | 300 |
| | | C_Ter8R | TCGTCAAGGGGAATATCAG | |
| | 9 | C_Ter9F | AACAATGTGGACAAGCACT | 300 |
| | | C_Ter9R | TACTGTGGGGGTCTAGTTG | |
| | 10 | C_Ter10F | CGGAATTACAGGTATCACAAC | 300 |
| | | C_Ter10R | CAAGATGTTCCTTTAACCAGTG | |

**Table S3.** PCR cycling conditions for DNA FISH probe production.

| Target | Primer | Product (nt) | Cycling conditions |
|---|---|---|---|
| *A. filiformis ori* | see Table S2 | Initial 3,622 nt-long fragment | 98°C for 5 min, followed by 31 cycles at 98°C for 20 s, 64°C (-0.3°C every cycle) for 20 s, 72°C for 120 s, followed by a final elongation step at 72°C for 5 min. |
| | see Table S2 | ~300 nt-long probes (P1-P11) | 98°C for 5 min, followed by 31 cycles at 98°C for 20 s, 64°C (-0.3°C every cycle) for 20 s, 72°C for 15 s, followed by a final elongation step at 72°C for 5 min. |
| *A. filiformis ter* | see Table S2 | Initial 3,583 nt-long fragment | 98°C for 5 min, followed by 31 cycles at 98°C for 20 s, 63°C (-0.3°C every cycle) for 20 s, 72°C for 120 s, followed by a final elongation step at 72°C for 5 min. |
| | see Table S2 | ~300 nt-long probes (P1-P11) | 98°C for 5 min, followed by 31 cycles at 98°C for 20 s, 62°C (-0.3°C every cycle) for 20 s, 72°C for 15 s, followed by a final elongation step at 72°C for 5 min. |
| *S. muelleri ori* | see Table S2 | Initial 8,158 nt-long fragment | 98°C for 5 min, followed by 31 cycles at 98°C for 20 s, 58°C (-0.3°C every cycle) for 20 s, 72°C for 240s, followed by a final elongation step at 72°C for 5 min. |
| | see Table S2 | ~300 nt-long probes (P1-P10) | 98°C for 5 min, followed by 31 cycles at 98°C for 20 s, 58°C (-0.3°C every cycle) for 20 s, 72°C for 15s, followed by a final elongation step at 72°C for 5 min. |
| *S. muelleri ter* | see Table S2 | Initial 8,187 nt-long fragment | 98°C for 5 min, followed by 31 cycles at 98°C for 20 s, 58°C (-0.3°C every cycle) for 20 s, 72°C for 240s, followed by a final elongation step at 72°C for 5 min. |

| | | | |
|---|---|---|---|
| | see Table S1 | ~300 nt-long probes (P1-P9) | 98°C for 5 min, followed by 31 cycles at 98°C for 20 s, 58°C (-0.3°C every cycle) for 20 s, 72°C for 15s, followed by a final elongation step at 72°C for 5 min. |
| *C. steedae* *ori* | see Table S2 | Initial 7,353 nt-long fragment | 95°C for 5 min, followed by 31 cycles at 95°C for 20 s, 58°C (-0.3°C every cycle) for 20 s, 72°C for 270s, followed by a final elongation step at 72°C for 5 min. |
| | see Table S2 | 300 nt-long probes (P1-P11) | 95°C for 5 min, followed by 31 cycles at 95°C for 20 s, 58°C (-0.3°C every cycle) for 20 s, 72°C for 30s, followed by a final elongation step at 72°C for 5 min. |
| *C. steedae* *ter* | see Table S2 | Initial 4, 498 nt-long fragment | 95°C for 5 min, followed by 31 cycles at 95°C for 20 s, 58°C (-0.3°C every cycle) for 20 s, 72°C for 270s, followed by a final elongation step at 72°C for 5 min. |
| | see Table S2 | 300 nt-long probes (P1-P10) | 95°C for 5 min, followed by 31 cycles at 95°C for 20 s, 58°C (-0.3°C every cycle) for 20 s, 72°C for 30s, followed by a final elongation step at 72°C for 5 min. |
| *Ca.* T. hypermnestrae *ori* | see Table S2 | Initial 2,858 nt-long fragment | 95°C for 5 min, followed by 31 cycles at 98°C for 45 s, 57°C for 45 s, 72°C for 180 s, followed by a final elongation step at 72°C for 5 min. |
| | see Table S2 | ~300 nt-long probes (P1-P9) | 95°C for 5 min, followed by 31 cycles at 98°C for 45 s, 57°C for 45 s, 72°C for 30 s, followed by a final elongation step at 72°C for 5 min. |
| *Ca.* T. hypermnestrae *ter* | see Table S2 | Initial 3,239 nt-long fragment | 98°C for 5 min, followed by 31 cycles at 98°C for 20 s, 63°C (-0.3°C every cycle) for 20 s, 72°C for 125 s, followed by a final elongation step at 72°C for 5 min. |

| | | | |
|---|---|---|---|
| | see Table S2 | ~300 nt-long probes (P1-P10) | 98°C for 5 min, followed by 31 cycles at 98°C for 20 s, 58°C for 20 s, 72°C for 15 s, followed by a final elongation step at 72°C for 5 min. |
| *Ca.* T. hypermnestrae<br>*ftsQAZ* | see Table S2 | Initial 4,010 nt-long fragment | 98°C for 5 min, followed by 31 cycles at 98°C for 20 s, 65°C (-0.3°C every cycle) for 20 s, 72°C for 120 s, followed by a final elongation step at 72°C for 5 min. |
| | see Table S2 | ~300 nt-long probes (P1-P11) | 95°C for 5 min, followed by 31 cycles at 94°C for 60 s, 57°C for 30 s, 72°C for 30 s, followed by a final elongation step at 72°C for 5 min. |
| *Ca.* T. hypermnestrae<br>*parS2* | see Table S2 | Initial 3,632 nt-long fragment | 95°C for 5 min, followed by 31 cycles at 98°C for 20 s, 64°C (-0.3°C every cycle) for 20 s, 72°C for 120 s, followed by a final elongation step at 72°C for 5 min. |
| | see Table S2 | ~300 nt-long probes (P1-P10) | 98°C for 5 min, followed by 31 cycles at 98°C for 20 s, 64°C (-0.2°C every cycle) for 20 s, 72°C for 20 s, followed by a final elongation step at 72°C for 5 min. |

**Table S4.** SRA accession numbers

| | **Sample name** | **NCBI accession number** |
|---|---|---|
| **Genome sequencing** | *A. filiformis* long reads | fast5: SRR13289602, fastq: SRR16893507 |
| | *A. filiformis* short reads | SRR15998661 |
| | *S. muelleri* long reads | fast5: SRR13289773, fastq: SRR16893579 |
| | *S. muelleri* short reads | SRR15998660 |
| | *C. steedae*, long reads | fast5: SRR16896741, fastq: SRR16947409 |
| | *C. steedae,* short reads | SRR19164917 |
| | *A. filiformis* assembly | GCF_020162295.1 |
| | *S. muelleri* assembly | GCF_020162275.1 |

| | | |
|---|---|---|
| | *C. steedae,* assembly | GCF_023547005.1 |
| **Marker frequency analysis** | *A. filiformis*, late exponential phase | SRR17498973 |
| | *A. filiformis*, stationary phase | SRR17498972 |
| | *S. muelleri*, late exponential phase | SRR17498978 |
| | *S. muelleri*, stationary phase | SRR17498977 |
| | *C. steedae*, late exponential phase | SRR17498976 |
| | *C. steedae*, stationary phase | SRR17498975 |
| | *E. coli,* exponential phase* | ERR902995 |
| | *E. coli,* stationary phase* | ERR902996 |
| | *A. filiformis* 1kb-binned raw counts | GSE194131 |
| | *S. muelleri* 1kb-binned raw counts | GSE194134 |
| | *C. steedae* 1kb-binned raw counts | GSE205139 |
| **RNA-Seq** | *A. filiformis*, exponential phase (3 biological replicates) | SRR17498971, SRR17498970, SRR17498961 |
| | *A. filiformis*, stationary phase (3 biological replicates) | SRR17498969, SRR17498968, SRR17498954 |
| | *A. filiformis*, plate (3 biological replicates) | SRR17498958, SRR17498959, SRR17498960 |
| | *S. muelleri*, stationary phase (3 biological replicates) | SRR17498936, SRR17498937, SRR17498935 |
| | *S. muelleri*, exponential phase (3 biological replicates) | SRR17498941, SRR17498942, SRR17498943, |
| | *S. muelleri*, plate (3 biological replicates) | SRR17498949, SRR17498940, SRR17498938 |
| | *A. filiformis* raw count table | GSE194132 |
| | *S. muelleri* raw count table | GSE194135 |
| | *C. steedae* raw count table | GSE250250 |
| | *C. steedae,* plate (3 biological replicates) | SRR17498948, SRR17498950, SRR17498939 |
| | *C. steedae,* exponential phase (3 biological replicates) | SRR17498951, SRR17498952, SRR17498953 |
| | *C. steedae,* stationary phase (3 biological replicates) | SRR17498945, SRR17498946, SRR17498947 |
| **3C-Seq** | *A. filiformis*, exponential phase (replicate 1) | SRR19164944 |
| | *A. filiformis*, exponential phase (replicate 2) | SRR19164943 |
| | *A. filiformis*, stationary phase | SRR19164942 |

| | | |
|---|---|---|
| | *A. filiformis*, plate (replicate 1) | SRR27205587 |
| | *A. filiformis*, plate (replicate 2) | SRR27206238 |
| | *A. filiformis*, exponential phase with 25 µg/ml Rifampicin for 30 min | SRR27206238 |
| | *A. filiformis*, exponential phase with 25 µg/ml Chloramphenicol for 30 min | SRR27206240 |
| | *S. muelleri*, plate (replicate 1) | SRR19164985 |
| | *S. muelleri*, plate (replicate 2) | SRR19164986 |
| | *S. muelleri*, exponential phase | SRR27206933 |
| | *S. muelleri*, stationary phase | SRR19164987 |
| | *S. muelleri*, exponential phase with 25 µg/ml Rifampicin for 30 min | SRR27206934 |
| | *S. muelleri*, exponential phase with 25 µg/ml Chloramphenicol for 30 min | SRR27206937 |
| | *C. steedae*, plate | SRR27206419 |
| | *C. steedae*, exponential phase | SRR27206418 |
| | *C. steedae*, stationary phase | SRR27206420 |
| | *A. filiformis* 3C-seq matrices | GSE250400 |
| | *S. muelleri* 3C-seq matrices | GSE250402 |
| | *C. steedae* 3C-seq matrices | GSE250401 |

* published by (Ivanova *et al.*, 2015)

**Table S5.** Genes situated in the looping regions in *A. filiformis* grown on solid medium. Only genes encoding for proteins with predicted function are listed. Genes are subdivided into eggNOG-based COG categories. n.a.: not available. See Supplementary Data Set 1 for details.

| *Cellular processing and signaling* | | |
|---|---|---|
| gene name | gene product | function |
| n.a. | adhesin | *host adhesion and/or immune evasion* |
| *Information storage and processing* | | |
| gene name | gene product | function |
| *tetR* | tetR family transcriptional repressor | *Transcription* |
| *Metabolism* | | |
| gene name | gene product | function |
| *dusA* | tRNA-dihydrouridine synthase A | *Coenzyme transport and metabolism* |

**Table S6.** Genes situated in the looping regions in *A. filiformis* grown in liquid culture. Only genes encoding for proteins with predicted function are listed. Genes are subdivided into eggNOG-based COG categories. n.a.: not available. See Supplementary Data Set 1 for details.

| *Information storage and processing* | | |
|---|---|---|
| gene name | gene product | function |
| *ruvC* | component of RuvABC resolvasome, endonuclease | *Replication, recombination and repair* |
| *prmC* | release factor glutamine methyltransferase | *Translation, ribosomal structure and biogenesis* |
| *Metabolism* | | |
| gene name | gene product | function |
| *lutA* | Lactate utilization protein A 1 | *Energy production and conversion* |
| *lldD* | L-lactate dehydrogenase, FMN linked | |
| *bioH* | Pimeloyl-[acyl-carrier protein] methyl ester esterase | *Lipid transport and metabolism* |
| *pncA* | Nicotinamidase/pyrazinamidase | *Secondary metabolites biosynthesis, transport and catabolism* |
